# Nanophysiology Approach Reveals Diversity in Calcium Microdomains across Zebrafish Retinal Bipolar Ribbon Synapses

**DOI:** 10.1101/2024.11.01.617078

**Authors:** Nirujan Rameshkumar, Abhishek P Shrestha, Johane M. Boff, Mrinalini Hoon, Victor Matveev, David Zenisek, Thirumalini Vaithianathan

**Affiliations:** Department of Pharmacology, Addiction Science, and Toxicology, The University of Tennessee Health Science Center, TN 38163; Department of Neuroscience, University of Wisconsin, Madison, WI, USA; McPherson Eye Research Institute, University of Wisconsin, Madison, WI, USA; Department of Ophthalmology and Visual Sciences, University of Wisconsin, Madison, WI, USA; Department of Mathematical Sciences, New Jersey Institute of Technology, Newark, NJ 07102, USA; Department of Cellular and Molecular Physiology, Yale University School of Medicine, New Haven, CT 06520-8066, USA; Department of Ophthalmology and Visual Sciences, Yale University School of Medicine, New Haven, CT 06520-8066, USA; Department of Ophthalmology, Hamilton Eye Institute, University of Tennessee Health Science Center, Memphis, TN 38163, USA

**Author notes:** These authors contributed equally to this work (Nirujan Rameshkumar, Abhishek P Shrestha, Johane M. Boff). Department of Surgery, College of Medicine, University of Florida, Gainesville, Florida (Abhishek P Shrestha). Contact: Thirumalini Vaithianathan, Ph.D.,.

## Abstract

Rapid and high local calcium (Ca^2+^) signals are essential for triggering neurotransmitter release from presynaptic terminals. In specialized bipolar ribbon synapses of the retina, these local Ca^2+^ signals control multiple processes, including the priming, docking, and translocation of vesicles on the ribbon before exocytosis, endocytosis, and the replenishment of release-ready vesicles to the fusion sites for sustained neurotransmission. However, our knowledge about Ca^2+^ signals along the axis of the ribbon active zone is limited. Here, we used fast confocal quantitative dual-color ratiometric line-scan imaging of a fluorescently labeled ribbon binding peptide and Ca^2+^ indicators to monitor the spatial and temporal aspects of Ca^2+^ transients of individual ribbon active zones in zebrafish retinal rod bipolar cells (RBCs). We observed that a Ca^2+^ transient elicited a much greater fluorescence amplitude when the Ca^2+^ indicator was conjugated to a ribeye-binding peptide than when using a soluble Ca^2+^ indicator, and the estimated Ca^2+^ levels at the ribbon active zone exceeded 26 μM in response to a 10-millisecond stimulus, as measured by a ribbon-bound low-affinity Ca^2+^ indicator. Our quantitative modeling of Ca^2+^ diffusion and buffering is consistent with this estimate and provides a detailed view of the spatiotemporal [Ca^2+^] dynamics near the ribbon. Importantly, our data demonstrates that the local Ca^2+^ levels may vary between ribbons of different RBCs and within the same cells. The variation in local Ca^2+^ signals is found to correlate with ribbon size and active zone extent. Our serial electron microscopy results provide new information about the heterogeneity in ribbon size, shape, and area of the ribbon in contact with the plasma membrane.

## Introduction

Sensory synapses in the retina rely on the proper function of a specialized organelle, the synaptic ribbon ^1–5^. Retinal bipolar cells serve as the major conduit for transmitting visual information across the vertebrate retina. Rod bipolar cells (RBCs) can release brief bursts of neurotransmitters to signal a change in contrast or sustain the continuous release of neurotransmitters in a graded manner to provide an analog read-out of luminance ^6^. To maintain this ability, the RBCs must exert dynamic control over neurotransmitter release rate and facilitate efficient recruitment of release-ready vesicles to fusion sites near the synaptic ribbon. It is established that the elevation of presynaptic Ca^2+^ in RBCs regulates both dynamic changes in the release rate and accelerates the rate of vesicle replacement ^6–14^. Of note, nanodomain calcium signals refer to highly localized, steep calcium concentration gradients that occur within tens to a couple of hundred nanometers of an open calcium channel. They are typically involved in triggering fast, synchronous neurotransmitter release, especially in synapses where Ca²⁺ sensors are closely coupled to the channels ^15–18^. Microdomain Ca^2+^ signals extend over a larger spatial range and arise when multiple calcium channels open in proximity, leading to overlapping calcium plumes. These domains result in slower, more diffuse Ca^2+^ elevations, often associated with modulatory functions or asynchronous release, where the Ca^2+^ sensor is located farther from the channel cluster ^18–20^. However, the spatiotemporal properties of the Ca^2+^ signals that control neurotransmitter release and the molecular entities that regulate the interplay between Ca^2+^ signals and synaptic vesicle dynamics to sustain kinetically distinct neurotransmitter release components remain poorly understood. Here, we begin to address this lack of knowledge by measuring local Ca^2+^ signals at positions along the synaptic ribbon at different distances from the active zone in retinal RBCs.

Resolving local Ca^2+^ signals is technically challenging as it requires information about the spatiotemporal properties of Ca^2+^ signals specific to ribbon sites. Previous studies of Ca^2+^ dynamics in goldfish and mammalian RBCs focused primarily on the terminal as a whole and used quantitative methods to examine bulk Ca^2+^ levels, which are significantly lower and slower than those occurring at the active zone ^21–24^. Our previous studies estimated that the Ca^2+^ signals at a single ribbon active zone in zebrafish RBCs likely exceed sub-micromolar concentration levels within 3 milliseconds after the opening of voltage-gated Ca^2+^ (Cav) channels located beneath the synaptic ribbon ^25^. However, these studies were based on estimates using soluble Ca^2+^ indicators to estimate the Ca^2+^ signals at the plasma membrane, which are free to diffuse and thus cause the spread of the signal. Measurements of Ca^2+^ signals along the axis of the ribbon have not been attempted, even though these signals likely control replenishment and priming of vesicles for slower phases of neurotransmitter release. A practical way to conceptualize the distinction between micro- and nanodomain Ca^2+^ signals is by approximating spatial scales: signals extending from around 0.5 μm and above may be considered microdomains, while those below 0.3-0.4 μm fall within the nanodomain range ^18^. To determine Ca^2+^ signals along the axis of zebrafish RBC ribbons, we establish a nanophysiology ratiometric approach that measures the spatial and temporal properties of Ca^2+^ transients by targeting the Ca^2+^ indicator to the ribbon via conjugation to a ribbon binding peptide (RBP)^26^. Resolving Ca^2+^ signals in the immediate vicinity of the active zone is not currently possible due to the finite spatiotemporal resolution of optical imaging. Thus, we employ a quantitative model using Ca^2+^ diffusion and buffering to resolve the RBC Ca^2+^ signals associated with the active zone and along the ribbon and provide a detailed description of the spatiotemporal Ca^2+^ dynamics across zebrafish RBC ribbons. Our nanophysiology approach for measuring local Ca^2+^ transients at a single ribbon with high spatiotemporal resolution provides the first evidence of heterogeneity of Ca^2+^ signals at zebrafish RBC ribbon synapses. Heterogeneity in local Ca²⁺ signals persisted in some ribbons even after the application of high concentrations of exogenous Ca²⁺ buffers, suggesting that the variability in the faster, smaller, and more spatially confined Ca²⁺ microdomains originates from Ca²⁺ influx through synaptic Ca²⁺ channel clusters at the base of the synaptic ribbon. Using serial block-face scanning electron microscopy (SBF-SEM), we found substantial variability in synaptic ribbon size, shape, and particularly the area of the ribbon adjacent to the plasma membrane in the zebrafish bipolar cell terminal. Thus, the observed heterogeneity of the Ca^2+^ microdomain is likely due to variability in the number of Ca^2+^ channels near each ribbon. Since local Ca^2+^ signals control kinetically distinct neurotransmitter release, heterogeneity in local Ca^2+^ signal may alter the rate of vesicle release and allow them to function independently, adding a new mechanism for increasing dynamic range of RBC.

## Results

### Ca^2+^ concentrations at single ribbon locations measured using high- and low-affinity diffusible indicators

Synaptic vesicles in RBCs are distributed among at least four distinct pools based on their fusion kinetics, which are assumed to reflect the average proximity of vesicles to Cav channels and the state of vesicle preparedness for Ca^2+^-triggered fusion ^1,6,8,11–13,27–31^ (see **Supplementary Fig. 1**). To visualize and measure Ca^2+^ signals near the RBC synaptic ribbon controlling such kinetically distinct components of neurotransmitter release, we established a quantitative nanophysiology approach shown in **Fig.1** (see also *Materials and Methods*). In this approach, both the ribbon location and the spatiotemporal Ca^2+^ signal profile were simultaneously measured by dialyzing the zebrafish RBC terminals with both TAMRA (tetramethyl rhodamine)-labeled RIBEYE-binding peptide (RBP) ^32^ (to label synaptic ribbons; **Fig. 1Aa**) and a high-affinity Ca^2+^ indicator Cal-520^®^ (Cal520HA, effective *K_D_* 795 nM; see *Materials and Methods*) using a whole-cell patch pipette placed directly at the cell terminal. A rapid x-t line scan was taken perpendicular to the plasma membrane across a ribbon, extending from the extracellular space to the cytoplasmic region beyond the ribbon (**Fig. 1Ab**). The spatial resolution was limited by the point spread function (PSF) of the microscope to approximately 270 nm (**Supplementary Fig.2A**). In our previous studies, line scans have been applied primarily to measure the temporal properties of Ca^2+^ signals ^25^. However, x-t raster plots obtained at a ribbon active zone labeled with RBP allow us to characterize the spatial localization of Ca^2+^ transients relative to the synaptic ribbon and plasma membrane (**Fig.1C-D**) in addition to characterizing its temporal aspects (**Fig.1E-F**). We previously used a similar approach for localizing and tracking single synaptic vesicles before and during fusion at a single ribbon ^33^, and to measure the kinetics of clearance of fused synaptic vesicle membrane in zebrafish RBC ^34^. We found that depolarization-evoked Ca^2+^ influx caused a rapid increase in Ca^2+^ signals at ribbon locations (**Fig.1C, cyan, white horizontal arrowhead**) and increased more slowly and less dramatically in the cytoplasm (**Fig.1C, cyan, white vertical arrow**) in zebrafish RBC. The Sigmoid-Gaussian function fitting of the x-t scans horizontal profile scans (see *Materials and Methods*) show that the local Ca^2+^ signals increased rapidly during stimuli (**Fig.1D**, cyan line) and then decreased immediately after the end of depolarization (**Fig.1D**, black line) approaching the spatial profile corresponding to the resting Ca^2+^ levels (**Fig.1D**, gray line). As expected, the centroid position of Ca^2+^ signals during depolarization (**Fig.1D, cyan** x_0_) is closer to the plasma membrane (**Fig.1D, magenta** x_1/2_) than the centroid position of RBP (**Fig.1D magenta**, x_0_), since Cav channels are located at the membrane ^25,32,35–48^.

**Fig 1.**
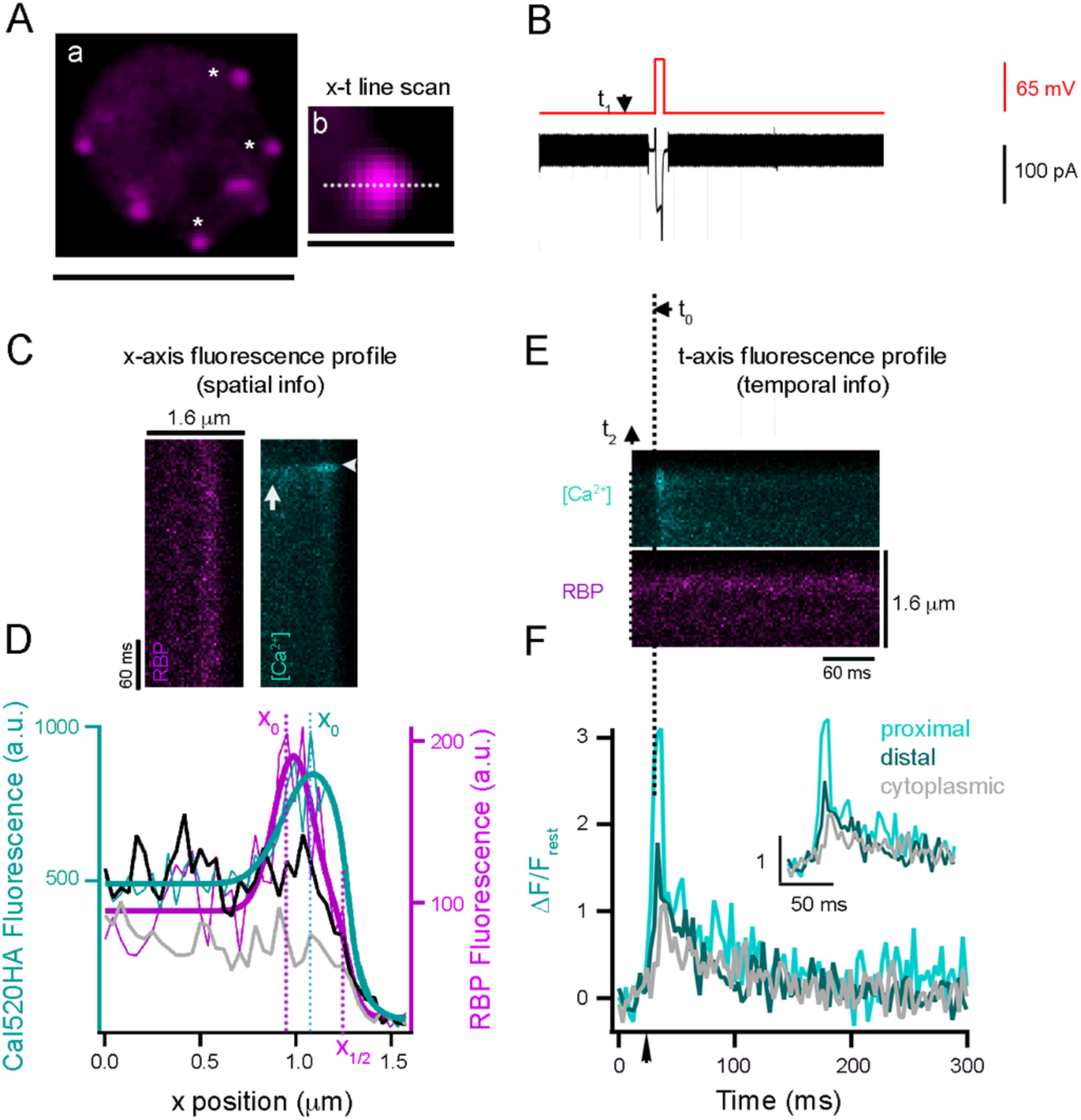
Nanophysiology approach unveils spatiotemporal properties of local Ca^2+^ signaling in retinal RBC terminals. **A. Left panel.** Single projection from a series of confocal optical sections through a zebrafish RBC synaptic terminal. A synaptic terminal was voltage-clamped using a whole-cell pipette with an internal solution containing TAMRA-RBP (magenta) to label synaptic ribbons (**a**, magenta). RBP fluorescence was concentrated at ribbons and also filled the entire terminal, allowing visualization of the terminal border. Experiments were carried out on ribbons that could be distinguished from adjacent ribbons (white asterisks). Scale, 5 μm. **Right panel**, Close-up view of a single synaptic ribbon. The outside of the cell is to the right, and x-t scan lines (dotted lines) were positioned perpendicular to the plasma membrane, extending from the intracellular side of the ribbon to the extracellular space. A rapid x-t line scan was taken at a ribbon location perpendicular to the plasma membrane across a ribbon (**b**) with sequential dual laser scanning performed at rates of 1.51 milliseconds per line per channel (3.02 milliseconds per line for both channels). The resulting x-t raster plots were used to measure the fluorescence intensity profiles of RBP (magenta) and the Ca^2+^ transient (cyan; Cal520HA) shown in panels **C** and **E**. Scale, 1.6 μm. **B.** Voltage-clamp recording of a RBC terminal. Terminals were held at -65 mV and stepped to 0 mV (t_0_) for 10 ms (red) to evoke a brief Ca^2+^ current (black). A typical experiment began with a voltage command (V_H_= -65 mV), and a TTL pulse generated by the Patch Master software (t_1_) triggers image acquisition (t_2_). t_0_ is the time of depolarization. **C.** Illustration of the approach used to obtain the spatial location of Ca^2+^ signals with respect to the ribbon. Example of an x-t raster plot that is oriented to illustrate the x-axis intensity profiles of RBP (magenta) and Ca^2+^ signal (cyan) fluorescence during a brief depolarization. Sequential dual laser scanning was performed at 1.51 milliseconds per line for one channel (3.02 milliseconds per line for both channels). **D.** Fluorescence intensity profiles along the x-axis for RBP (magenta) and Cal520HA before stimuli (gray line), during 10 ms depolarization (cyan line), and after stimuli (black line) depolarization were obtained by averaging three pixels along the time-axis. RBP (magenta) and Ca^2+^ signals during (light cyan) were fit with a Sigmoid-Gaussian function ^1^. The centroid (x-axis position) of the RBP (magenta) and Ca^2+^ signals during (cyan) were taken as the peak of the Gaussian fit (x_0_). The parameter x_1/2_ (dotted magenta line) from the Sigmoid fit to the RBP fluorescence (magenta trace) was used to estimate the location of the plasma membrane. **E.** The x-t raster plot shown was from the same recording as in **panel C** but re-oriented to demonstrate the t-axis intensity profiles of RBP (magenta) and Ca^2+^ transient (cyan). **F.** Spatially averaged Cal520HA fluorescence as a function of time at ribbon proximal (light cyan), distal (dark cyan), and cytoplasmic (gray) locations from the single ribbon shown in **D**, upper panel.

To characterize the temporal Ca^2+^ profile at different distances from the plasma membrane, we analyzed the x-t line scans along the time-axis at three distinct distances (**Fig. 1E-F**) based on the spatial profile of the ribbon described in **Fig.1C-D** (see also *Materials and Methods*). We will refer to the corresponding three measurements as *ribbon-proximal*, *ribbon-distal*, and *cytoplasmic* (see *Materials and Methods*). We found that at the onset of the stimulus (**Fig. 1F**, black arrow), the fluorescence at the location near the ribbon proximal to the membrane (ribbon-proximal) rose more rapidly and to a higher level (**Fig. 1F**, light cyan trace) than that at the ribbon-distal and cytoplasmic locations (**Fig. 1F**, dark cyan and gray traces, respectively). To quantify the kinetics of ribbon-proximal and distal Ca²⁺ signals, we averaged multiple x-t scans acquired with Cal520HA under the same imaging conditions (**Fig. 2A**). The maximum values of trial-averaged Δ*F*/*F*_rest_ (changes in the Cal520HA fluorescence during depolarization normalized to background level before depolarization) were significantly higher at ribbon-proximal locations (nearest the Cav channels) than those at ribbon-distal locations, with mean ± SEM of 1.9 ± 0.3 and 1.5 ± 0.25, respectively (**Fig. 2A**; p< 0.001, N=24). These findings suggest that the spatial resolution of our Ca^2+^ imaging using Cal520HA is sufficient to resolve differences in the smaller Ca^2+^ signals at ribbon proximal and distal locations. However, it should be noted that Cal520HA will be partially saturated at the Ca^2+^ levels expected in Ca^2+^ microdomains relevant for vesicle exocytosis ^49^, affecting both the amplitude and kinetics of the fluorescence signal. Therefore, we repeated the x-t line scan analysis with a lower-affinity soluble Ca^2+^ indicator Cal520LA (*K_D_* 90 μM; **Fig. 2B**) allowing us to better define the typical Ca^2+^ signals controlling distinct neurotransmitter release components corresponding to locations proximal and distal relative to the synaptic ribbon (**Fig. 2B**). We found that the ribbon-proximal signals detected with Cal520LA (**Fig. 2B**, light cyan) showed a sharper decay at the termination of the stimulus (**Fig. 2B**), when compared to ribbon-distal signals (**Fig. 2B**, dark cyan), as one would expect for nanodomain Ca^2+^ elevations ^50–52^. In twenty-one similar experiments, the peak Δ*F*/*F*_rest_ at the membrane after 10-ms depolarization was significantly larger for proximal than distal signals (Δ*F*/*F*_rest_: 3.1 ± 0.4 and 1.9 ± 0.2 respectively P=0.001, N= 21). As expected, the ratio of proximal to distal signals measured with Cal520LA (1.6) was significantly higher than that measured with Cal520HA (1.3). The decay phase of all fluorescence transients was fit by a sum of two exponential functions, as described in Methods; for the Cal520HA recording (**Fig. 2A**) the two decay time constants and their relative magnitudes were similar for the ribbon-proximal and distal recordings, namely 11ms (60%) and 203ms (40%) for the ribbon-proximal recording, vs. 16ms (61%) and 208ms (39%) for the distal site. For the low affinity Cal520LA dye (**Fig. 2B**), the fluorescence decay components appeared faster due to faster Ca^2+^ unbinding from the dye, at 9.3ms (69%) and 175ms (31%) for the ribbon-proximal location, vs. 9ms (41%) and 135ms (59%) for the distal location. We note, however, that bi-exponential data fits are known to be highly sensitive to measurement duration and noise. Therefore, precise quantitative conclusions should not be drawn from the best-fit decay time constant values.

**Fig. 2.**
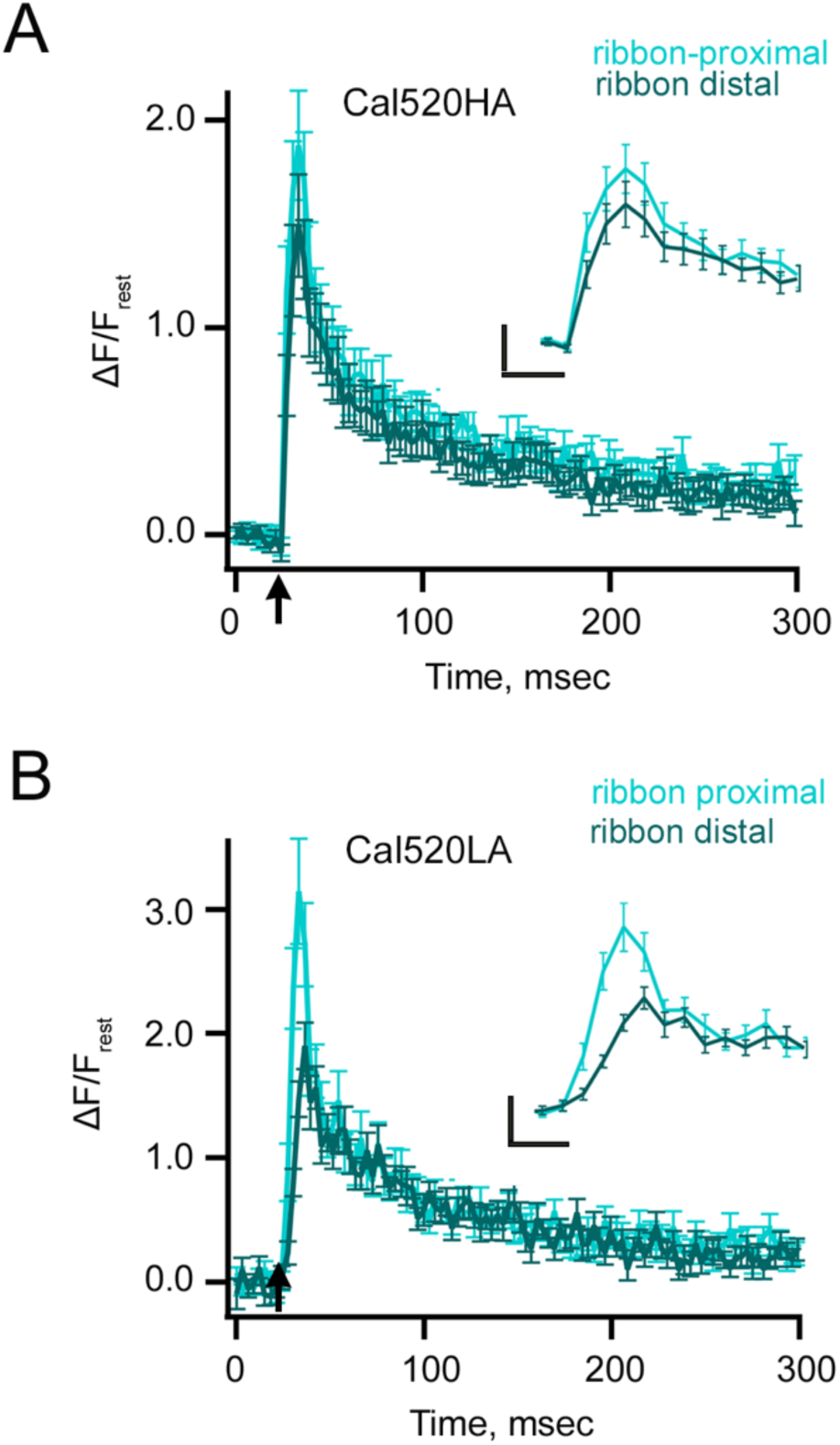
Kinetics of Ca^2+^ transients in response to brief stimuli recorded with freely diffusible indicators. **A.** Spatially averaged Cal520HA fluorescence as a function of time at ribbon proximal (light cyan) and distal (dark cyan) locations from a single ribbon, as shown in **Fig.1** (n=24 ribbons from 7 different RBCs). The corresponding maximum values of trial-averaged Δ*F*/*F*_rest_ were significantly higher at ribbon-proximal locations than those at ribbon-distal locations, with mean ± SEM of 1.9 ± 0.3 and 1.5 ± 0.25, respectively (paired t-test, p< 0.001, N=24). **A inset.** The temporal profile of events between 20-60 ms is shown in an expanded view for better visualization. Scale bars: vertical, 0.5 (ΔF/F_rest_); horizontal,10 ms. **B.** Spatially averaged Cal520LA fluorescence as a function of time at ribbon proximal (light cyan) and distal (dark cyan) locations. Data points show the average intensity (± SEM) in each horizontal row of 5 pixels for three 10 ms depolarizations at distinct ribbon locations (see *Materials and Methods* and **Fig.1**). Fluorescence intensity is normalized with respect to the baseline fluorescence before stimulation, and averaged over all pixels (i.e., over space and time). The arrow indicates the onset of the 10-ms depolarizing stimulus. (n=21 ribbons from 4 different RBCs). The peak Δ*F*/*F*_rest_ at the membrane after 10-ms depolarization was significantly larger for proximal than distal signals (paired t-test, Δ*F*/*F*_rest_: 3.1 ± 0.4 and 1.9 ± 0.2, respectively, P=0.001, N=21). The currents were not significantly different between the Cal520HA and Cal520LA conditions (unpaired t-test, Cal520HA average current = 45.1 ± 4.5 pA; Cal520LA average current = 51.4 ± 4.4 pA; p = 0.33). **B *inset*.** Temporal profile of events between 20-60 ms was expanded for better visualization. Scale bars: vertical, 0.5 (ΔF/F_rest_); horizontal, 10 ms.

### Ca^2+^ concentrations at single ribbon locations measured using ribbon-bound indicators

Although Ca²⁺-sensitive fluorescent chemical dyes have been used previously ^7,26,35,42,48,49,51–63^, the visualization of signals within smaller domains using freely diffusible Ca²⁺ reporters is limited by the resolution of light microscopy and the spread of the indicator by diffusion ^64^. Diffusible Ca^2+^ indicators report space-averaged Ca^2+^ concentrations, and their intracellular diffusion inherently broadens the spatial resolution of Ca^2+^ nanodomains. To partially overcome this problem, we targeted the Ca^2+^ indicators to the ribbon by fusing them to the RBP, as described previously ^26^. RBP-conjugated Ca^2+^ indicator s Cal520HA-RBP or Cal520LA-RBP were introduced to the RBC terminal together with fluorescently labeled RBP via whole-cell voltage clamp by placing the patch pipette directly at the terminal while imaging the terminal using laser scanning confocal microscopy. For two-color imaging, ribbons were labeled with TAMRA-RBP that did not interfere with Cal520-RBP fluorescence, and both channels were scanned sequentially to prevent possible bleed-through. Under these conditions, we used the spots detected by TAMRA-RBP to define the locus of the synaptic ribbon (**Fig 3A&B, magenta**). We also found punctate regions with both Cal520HA-RBP (**Fig 3A, cyan**) and Cal520LA-RBP (**Fig 3B, cyan**) at the same location, on a dimmer fluorescent background of the synaptic terminal, which correspond to the Ca^2+^ indicator-peptide complexes that are bound and not bound to the ribbon, respectively ^26^. The overall changes in Δ*F*/*F*_rest_ were averaged over several trials for proximal and distal Ca^2+^ signals measured with Cal520HA-RBP (**Fig. 3C**, light cyan vs. dark cyan) and Cal520LA-RBP (**Fig. 3D**, light cyan vs. dark cyan). When compared to their distal location counterparts, the Ca^2+^ signals proximal to the membrane showed a sharper decay at the termination of stimuli (**Figs. 3C** and **3D**, light vs. dark cyan trace), as expected when comparing nano- and microdomain Ca^2+^ profiles. Our data shows that the amplitude differences between ribbon-proximal and ribbon-distal Ca^2+^ signals were well resolvable using the ribbon-bound Cal520HA-RBP indicator (**Fig. 3C**, light vs. dark cyan traces: Δ*F*/*F*_rest_ = 3 ± 0.4 vs. 1.9 ± 0.3, respectively, p=0.001) and Cal520LA-RBP indicator (**Fig. 3D**, light vs. dark cyan traces: Δ*F*/*F*_rest_ = 5.5 ± 0.9 vs. 3.3 ± 0.8, respectively, p=0.003). The amplitudes of ribbon-proximal Ca^2+^ signals were higher when measured with Cal520LA-RBP than with Cal520LA-free (**Fig. 3E**, Cal520LA-RBP (light cyan) vs. Cal520LA-free (gray): Δ*F*/*F*_rest_ = 5.5 ± 0.9, n=30 vs. 3 ± 0.4, n=19, p=0.04) but this was not the case for distal Ca^2+^ signals (**Fig. 3F**, Cal520LA-RBP (light cyan) vs. Cal520LA-free (gray): Δ*F*/*F*_rest_ = 3.3 ± 0.8, *n*=30 vs. 1.9 ± 0.2, *n*=21, *p*=0.15). Notably, the Ca^2+^ signals at distal sites measured with Cal520LA-RBP reached their peak amplitude earlier (**Fig. 3F *inset***, dark cyan) than those measured with Cal520LA-free (**Fig. 3F *inset***, light gray). The two decay time constants and their relative magnitudes for the Cal520HA-RBP fluorescence (**Fig. 3C**) were 11ms (52.5%) and 182ms (47.5%) for the ribbon-proximal site, vs. 24ms (47%) and 192ms (53%) for the site distal to the ribbon. For the low affinity ribbon-bound indicator dye (**Fig. 3D**), the fluorescence decay components appeared faster, at 5.7ms (83%) and 129ms (17%) for the proximal site, vs. 7.5ms (76%) and 158ms (24%) for the distal location. Together, these findings suggest that conjugating the Cal520LA indicator to RBP provides a more accurate, promising approach for measuring the distinct local Ca^2+^ signals at ribbon-proximal vs. ribbon-distal locations.

**Fig.3.**
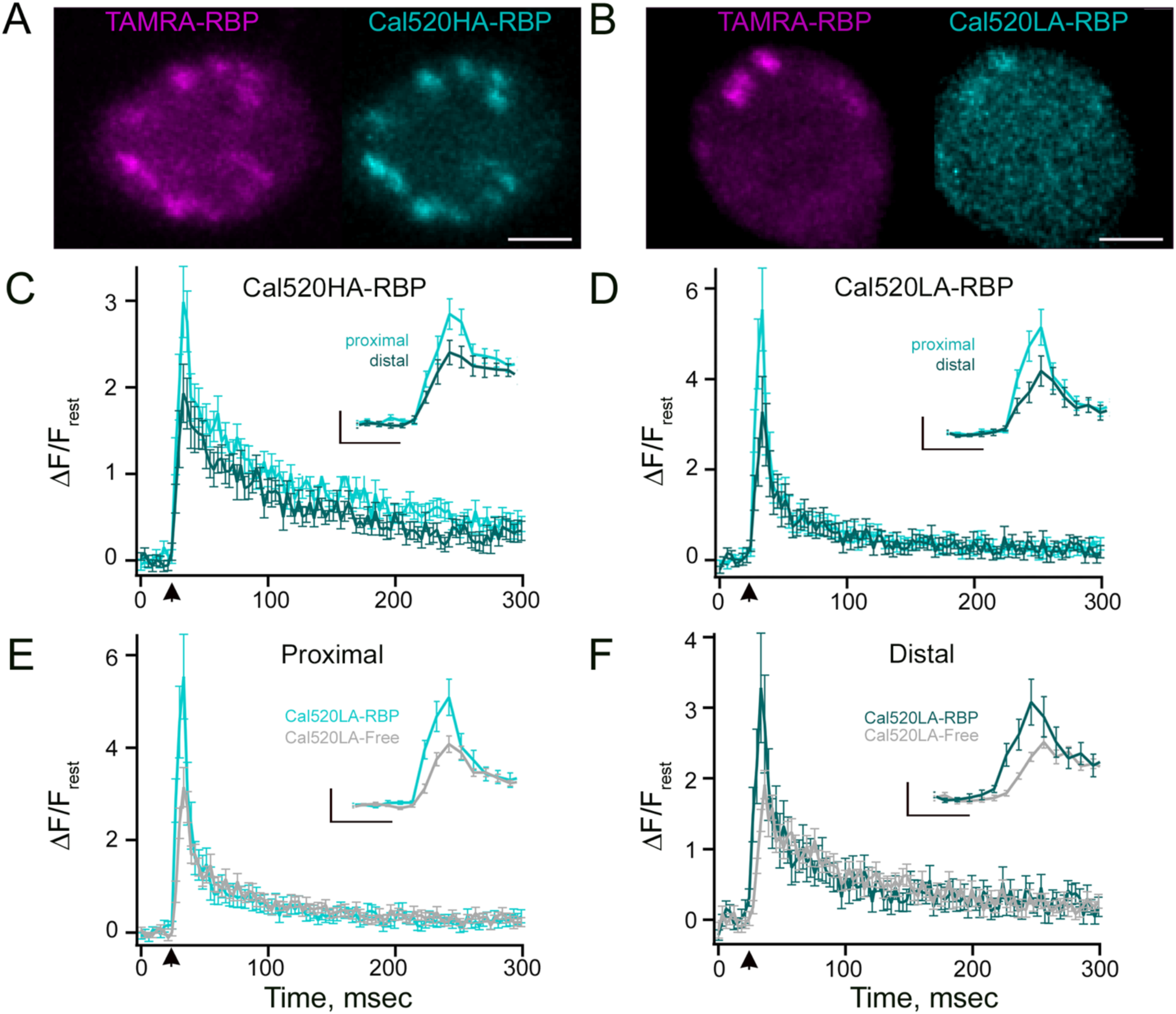
Temporal properties of Ca^2+^ transients recorded with free and ribeye-bound Ca^2+^ indicators. **A-B.** Confocal images of the isolated RBCs that were whole-cell voltage-clamped using an internal solution containing the TAMRA-RBP (magenta) and either (**A**) Cal520HA-RBP (cyan) or (**B**) Cal520LA-RBP (cyan). Note prominent spots in both TAMRA-RBP and ribeye-bound Ca^2+^ indicators (**A**) Cal520HA-RBP (cyan) or (**B**) Cal520LA-RBP, showing the location of the ribbon. Scale bars, 2 μm. **C-D.** Spatially averaged fluorescence of (**C**) Cal520HA-RBP (n=19) or (**D**) Cal520LA-RBP (n=30) as a function of time at ribbon proximal (light cyan), and distal (dark cyan) locations. Data points show the average intensity (± SEM) in each horizontal row of five pixels for 10 ms depolarizations at distinct ribbon locations. Fluorescence intensity at the onset of the 10 ms depolarizing stimulus (arrow) was normalized to the baseline fluorescence before stimulation and averaged over all pixels (i.e., over space and time). The amplitude differences between ribbon-proximal and ribbon-distal Ca^2+^ signals were well-resolvable using the ribbon-bound Cal520HA-RBP indicator (**Fig. 3C**, light vs. dark cyan traces, paired t-test: Δ*F*/*F*_rest_ = 3.0 ± 0.4 vs. 1.9 ± 0.3, respectively, p=0.001) and Cal520LA-RBP indicator (**Fig. 3D**, light vs. dark cyan traces, paired t-test: Δ*F*/*F*_rest_ = 5.5 ± 0.9 vs. 3.3 ± 0.8, respectively, p=0.003). The current amplitudes were not significantly different between Cal520HA-RBP and Cal520LA-RBP readings (mean current amplitudes: Cal520HA-RBP = 49.8 ± 2.2 pA, Cal520LA-RBP = 49.9 ± 2.9 pA; p = 0.99). **C-D inset.** Temporal profile of events between 10-50 ms were expanded for better visualization. Scale bars: vertical, 1 (ΔF/F_rest_, **C inset**) or 2 (ΔF/F_rest_, **D inset**); horizontal, 10 ms. **E-F.** Average fluorescence intensity of (**E**) proximal and (**F**) distal Ca^2+^ signals obtained with Cal520LA-RBP (light cyan and dark cyan, respectively) and Cal520L-free (grey). The amplitudes of ribbon-proximal Ca^2+^ signals were higher when measured with Cal520LA-RBP than with Cal520LA-free (**Fig. 3E**, Cal520LA-RBP (light cyan) vs. Cal520LA-free (gray), unpaired t-test: Δ*F*/*F*_rest_ = 5.5 ± 0.9, n=30 vs. 3.0 ± 0.4, n=19, p=0.04) but this was not the case for distal Ca^2+^ signals (**Fig. 3F**, Cal520LA-RBP (light cyan) vs. Cal520LA-free (gray), unpaired t-test: Δ*F*/*F*_rest_ = 3.3 ± 0.8, *n*=30 vs. 1.9 ± 0.2, *n*=21, *p*=0.15). **E-F *inset*.** Events between 10-50 ms were expanded for better visualization. Scale bars: vertical, 2 (ΔF/F_rest_, **E inset**) and 1 (ΔF/F_rest_, **F inset**); horizontal, 10 ms.

This method also enables ratiometric measurements with RBP-conjugated Ca^2+^ indicator by normalizing its fluorescence to that of RBP-peptides conjugated to Ca^2+^-insensitive fluorophores (TAMRA-RBP) to provide an estimate of Ca^2+^ concentration in RBC synaptic ribbons. We obtained the effective *K_1/2_* (*K*_eff_) by measuring the Cal520HA-RBP/ TAMRA-RBP fluorescence ratio in buffered Ca^2+^ solutions and using the Grynkiewicz equation^59^. For Cal520HA-RBP, we found the *K*_eff_ to be ∼795 nM, which is higher than the value of 320 nM reported by the manufacturer for Cal520HA. The differences between the in-cell measurements and the manufacturer’s values are likely to arise from differences in the cellular buffering capabilities, changes in dye properties due to interactions with the molecules inside the cell ^42,65–67^, and possible differences in the binding properties of the peptide-conjugated Ca^2+^ indicators when bound to synaptic ribbons. Since the in-cell approach most closely matched the experimental conditions, we used this value (*K*_eff_) for all further calculations.

We first measured the ribbon-proximal and ribbon-distal Ca^2+^ concentrations using Cal520HA-RBP with 0.2 mM EGTA in the patch pipette, as it allowed to resolve the gradient between the two signals (**Fig. 1A**). Under these conditions, we found a maximum ribbon-proximal Ca^2+^ concentration produced in response to a brief 10- ms pulse of 3.7 µM, and a maximum ribbon-distal Ca^2+^ concentration of 0.7 µM. These *apparent* Ca^2+^ concentration amplitudes are well below levels required to trigger exocytosis of the ultrafast releasable pool (UFRP) and readily releasable pool (RRP) ^49^ and, as discussed earlier, likely represent the lower bounds of the Ca^2+^ concentration at a single ribbon location due to local saturation of the high-affinity indicator and/or due to nanodomains that are smaller than the resolution attainable using light microscopy. It should also be noted that unlike freely diffusing Ca^2+^ indicators, RBP-conjugated indicators that slowly unbind from the ribbon are not readily replaced by free Ca^2+^ indicators, rendering them even more prone to saturation. Nevertheless, our data demonstrate that ribbon-bound indicators may better report local Ca^2+^ concentrations due to their localization, albeit still subject to the limitations of light microscopy.

To test the contribution of local saturation and to better estimate ribbon Ca^2+^ concentrations, we repeated our measurements using the low-affinity Ca^2+^ indicator Cal520LA conjugated to the RBP peptide. Because it was difficult to perform in-cell measurements to determine the *K*_eff_ for Cal520LA-RBP given the large amounts of Ca^2+^ required to calibrate the Cal 520LA indicator (see *Methods and Materials*), we used the *K_1/2_ value* provided by the manufacturer for free Cal520LA (*K_D_* 90 µM). However, we expect that in-cell measurements of *K*_eff,_ are likely to be different due to the cellular buffering properties reported previously for *K*_eff,_ measurements of Oregon Green BAPTA-5N in inner hair cells ^42^.

Bipolar cells release neurotransmitters primarily from ribbon active zones, although some release also occurs at non-ribbon sites (referred to as NR, **Fig. 4A**) ^68–70^. To reveal the Ca^2+^ signaling at and away from the ribbon, we performed whole-cell patch clamping and x-t line scans at ribbon (**Fig. 4Ba**, R) and non-ribbon (**Fig. 4Bb**, NR) sites perpendicular to the plasma membrane using TAMRA-RBP and Cal520LA-RBP. A brief (10-ms) depolarizing voltage-clamp pulse evoked rapid high Ca^2+^ signals (**Fig. 4Ba**, cyan raster plot) at ribbon locations (**Fig. 4Ba**, magenta raster plot) but not at non-ribbon locations (**Fig. 4Bb**, cyan raster plot). The amplitude of Ca^2+^ signals elicited by brief depolarization and detected with Cal520LA-RBP at the ribbon-proximal site (**Fig. 4C**, light cyan traces) were significantly higher than those at ribbon-distal (**Fig. 4C**, dark cyan traces) and non-ribbon (**Fig. 4C**, blue traces) sites. The Ca^2+^ concentration gradient along the ribbon is summarized in **Fig. 4D**. We found that the average Ca^2+^ concentration at the proximal side of the ribbon (26.4 ± 3.1 µM, n=26) was significantly different from that at the ribbon-distal (15.6 ± 1.5 µM, n=26) and non-ribbon (10.4 ± 0.4 µM, n=15) sites, and that Ca^2+^ concentrations at ribbon-distal sites is higher than those at non-ribbon sites (**Fig. 4D**). These measurements display large heterogeneity across distinct ribbons and distinct cells, with the coefficient of variation of about 60% for [Ca^2+^] measurements at locations proximal to the ribbon, compared to a 10% CV for multiple depolarizations for the same ribbon. The Ca^2+^ signals at distal sites also reached their peak amplitude earlier (**Fig. 4C *inset***, dark cyan arrowhead) than those at the non-ribbon sites (**Fig. 4C *inset***, blue arrowhead). These findings are consistent with non-ribbon vesicle release being governed by cytoplasmic residual Ca^2+^ or Ca^2+^ influx via clustered Cav channels at ribbon sites rather than non-ribbon active zones with clusters of Cav channels and are also in agreement with previous reports regarding the temporal delay and sensitivity of non-ribbon exocytosis to EGTA ^69,71^.

**Fig.4.**
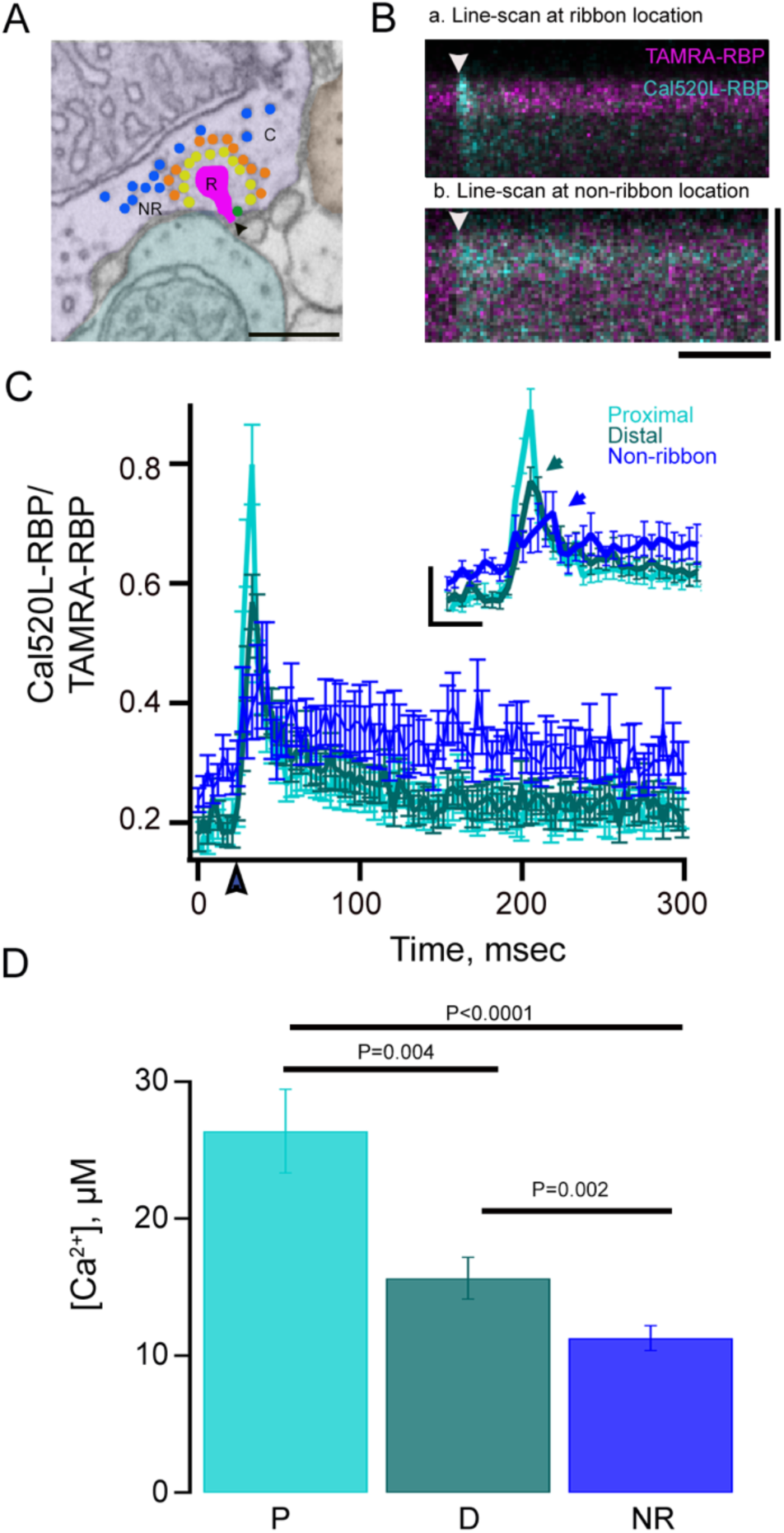
Ca2+ signals at synaptic ribbon at different distances from the plasma membrane. **A.** Ultrastructure of a zebrafish RBC with kinetically distinct vesicle pools, as described in **Supplementary Fig. 1**, UFRP (green vesicles), and RRP (yellow vesicles) are primarily released via ribbon sites (R) at the cytomatrix of the active zone (arrowhead). Recycling pool (RP, orange vesicles) in the cytoplasm (C), likely to be released via non-ribbon (NR) sites. Scale bar: 500 nm. **B.** Representative *x*-*t* plots show the fluorescence intensity of Cal520LA-RBP (cyan) and TAMRA-RBP (magenta) as a function of distance (vertical axis) and time (horizontal axis) at (**Ba**) ribbon sites and (**Bb**) non-ribbon (NR) locations. The darker region at the upper edge of each plot is the extracellular space and the arrowheads show the timing of depolarization. Scale bars: vertical, 1.6 μm and horizontal, 75 ms. **C.** Spatially averaged Cal520LA-RBP as a function of time at ribbon proximal (light cyan), distal (dark cyan), and non-ribbon locations (blue) (n=26 (proximal ribbons), 26 (distal ribbons), and 15 (non-ribbon) from 5∼8 different RBCs, respectively). **C *inset***. Temporal profile of events between 0-100 ms were expanded for better visualization. Scale bars: vertical, 0.2 (ΔF/F_rest_); horizontal, 20 ms. **D.** Ca^2+^ measurements along the ribbon axis using the nanophysiology approach demonstrate that the proximal Ca^2+^ signals can go as high as 26.4 ± 3.1 µM (light cyan, N=26) and distal as 15.6 ± 1.5 µM (dark cyan, N=26), and non-ribbon 10.4 ± 0.4 µM (NR, blue, N=15), respectively, in response to 10 ms stimuli. Error bars show standard errors. All conditions were significantly different from each other as assessed by paired t-test when comparing proximal vs. Distal and unpaired t-test when comparing non-ribbon to proximal or distal (proximal vs. distal p = 0.004; proximal vs. non-ribbon p < 0.001; distal vs. non-ribbon p = 0.002). The currents were not significantly different between conditions (Mean current: 0.2 mM EGTA Cal520L-RBP proximal and distal = 51.8 ± 3.2 pA, 0.2 mM EGTA Cal520L-RBP non-ribbon = 47.3 ± 3.0 pA; p = 0.35).

### Sensitivity of microdomain Ca^2+^ to exogenous buffers

Previous work has shown that the rate of vesicle replacement in RBCs is accelerated by elevated Ca^2+^ levels at sites along the ribbon, and that while millimolar levels of EGTA have little effect on the fast exocytosis component near Cav channels, they selectively block sustained exocytosis, likely by preventing Ca^2+^ from reaching the distal locations on the ribbon ^11,72–75^. These findings raise the possibility that Ca^2+^ has two sites of action, one near the Cav channels that trigger vesicle release and one further away that replenishes the supply of releasable vesicles. Burrone *et al.* (2002) proposed that endogenous Ca^2+^ buffers regulate the size of the RRP by limiting the spatial spread of Ca^2+^ ions and could suppress vesicle release at the periphery of the active zone in bipolar cells ^7^. However, the Ca^2+^ gradient at different locations along the ribbon controlling RRP release has never been examined. Thus, we used our higher resolution approach (**Fig.1**) to measure the Ca^2+^ gradient along the synaptic ribbon under buffering conditions that differentially modulate vesicle release and resupply. To determine the spatiotemporal properties of Ca^2+^ signals under these conditions, we performed rapid x-t line scans at a single ribbon location in the presence of ribbon-bound Cal520LA-RBP and under varying concentrations of exogenous buffer in the patch pipette solution, including (EGTA at 0.2 (**Fig. 5A**), 2 (**Fig. 5B**), and 10 mM (**Fig. 5C**) or BAPTA at 2 mM (**Fig. 5D**). In these experiments, we chose Cal520LA-RBP to study the varying contributions of exogenous buffer, as Cal530HA-RBP measurements are less accurate. However, it should be noted that for briefer depolarizations and lower amplitude depolarizations, where the signal for the Cal520LA-RBP is small, the Cal520HA-RBP could be more useful. We found that the ratio between proximal and distal Ca^2+^ signal amplitudes was similar in 0.2 and 2 mM EGTA (proximal-to-distal ratio ∼1.7, **Supplementary Table 1**) but was enhanced with 10 mM EGTA (proximal-to-distal ratio ∼1.9, **Supplementary Table 1**) and further enhanced with 2 mM BAPTA (proximal-to-distal ratio ∼4.5, **Supplementary Table 1**). These findings once again emphasize the reliable measurement of resolving ribbon-proximal vs. ribbon distal Ca^2+^ signals using our nano-physiology approach. Our experimental findings of the increase in the proximal-to-distal Ca^2+^ concentration ratio with increasing EGTA concentration are consistent with our simulation results (**Fig. 7**), although the corresponding ratios are greater in the simulation than in the experiment due to the large size of the microscope’s point-spread function.

**Fig. 5.**
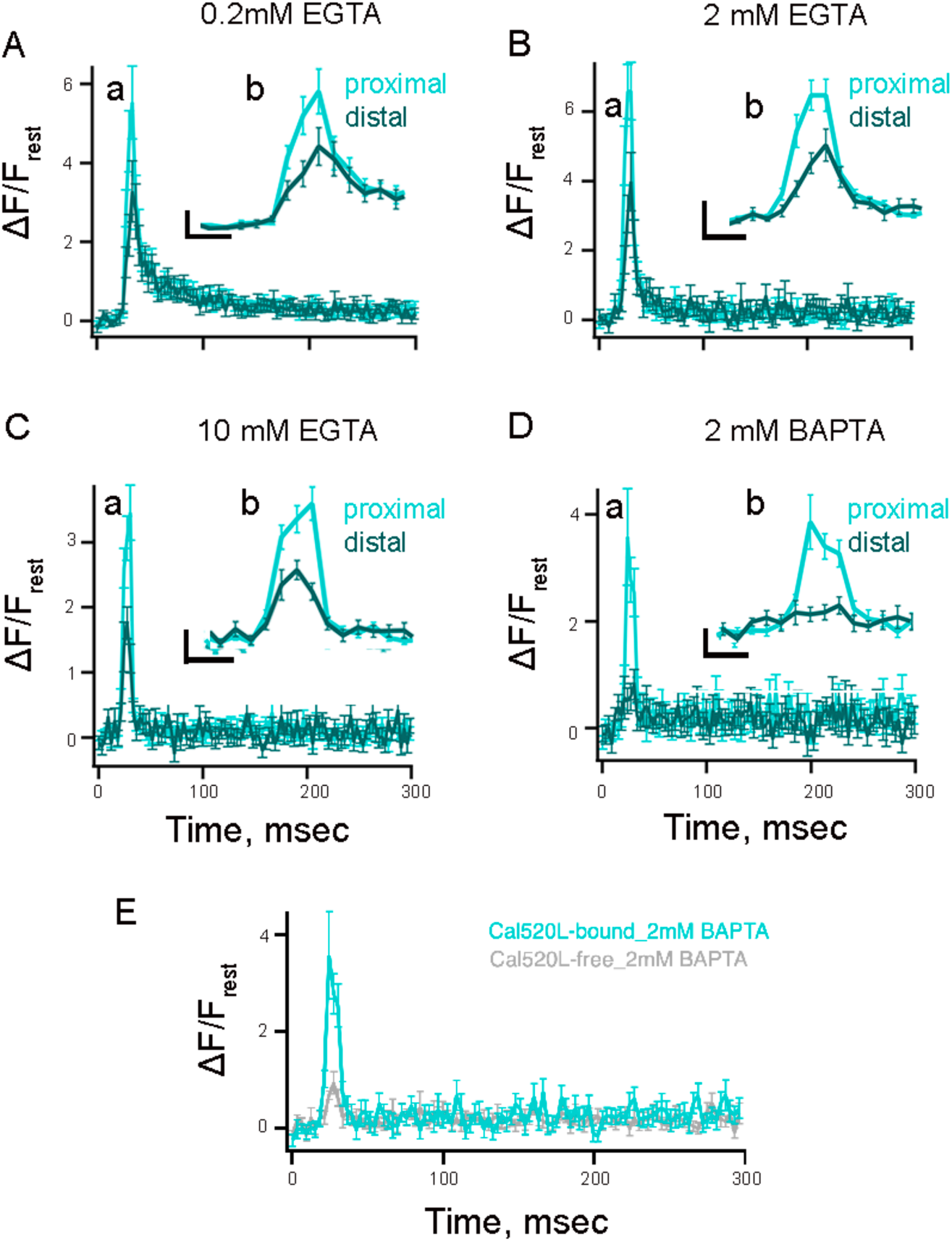
Effect of exogenous Ca^2+^ buffers on spatiotemporal properties of Ca^2+^ microdomains in RBC terminal recorded with low-affinity ribbon-bound dye. **A-D.** Average temporal fluorescence intensity (normalized to ΔF/F_rest_) of proximal (light cyan) and distal (dark cyan) Ca^2+^ signals with Cal520LA-RBP as a function of time at distinct ribbon locations with pipette solution containing (**A**) 0.2 mM EGTA, (**B**) 2 mM EGTA, (**C**) 10 mM EGTA, or (**D**) 2 mM BAPTA. Proximal measurements were significantly higher than distal measurements in all conditions as assessed using paired t-tests (0.2 mM EGTA: p = 0.0027, n = 29; 2 mM EGTA: p = 0.034, n = 21; 10 mM EGTA: p < 0.001, n = 42; 2 mM BAPTA: p = 0.0073, n = 21). The currents between conditions were not significantly different from each other (mean current amplitudes in 0.2 mM EGTA: 50.6 ± 3.0 pA, 2 mM EGTA: 49.7 ± 3.1 pA, 10 mM EGTA: 47.3 ± 3.1 pA, 2 mM BAPTA: 56.1 ± 2.6 pA; 0.2 mM EGTA vs 2 mM EGTA: p = 0.84, 0.2 mM EGTA vs 10 mM EGTA: p = 0.46, 0.2 mM EGTA vs 2 mM BAPTA: p = 0.19). **Inset.** The temporal profiles of events between 0-50 ms were expanded for better visualization. Scale bars: vertical, 2 (ΔF/F_rest_; **panels A and B**) and 1 (ΔF/F_rest_; **panels C and D**); horizontal, 10 ms. **E.** Average temporal fluorescence intensity (normalized to ΔF/F_rest_) of proximal Ca^2+^ signals measured with Cal520LA-RBP (cyan) and Cal520LA-free (gray) as a function of time with pipette solution containing 2 mM BAPTA. The corresponding maximum values of trial-averaged Δ*F*/*F*_rest_ was significantly higher with 2mM Cal520L bound BAPTA than 2 mM Cal520L free BAPTA, with mean ± SEM of 3.6 ± 0.9 and 1.3 ± 0.2, respectively (paired t-test, p = 0.015; Cal520L-free: n = 20; Cal520L-bound: n = 21). The currents between conditions were not significantly different from each other (mean current amplitudes in 2 mM BAPTA Cal520L-RBP: 56.1 ± 2.6 pA; 2 mM BAPTA Cal520L-Free: 53.3 ± 2.2 pA; p = 0.42).

Though BAPTA significantly abolished the spread of Ca^2+^ signals, it is impressive that a substantial amount of ribbon-proximal Ca^2+^ concentration measured with ribbon-bound indicators was still present with 2 mM BAPTA. One reason for this observation could be that ribbon-bound Ca^2+^ indicators are likely to measure Ca^2+^ signals very close to Cav channels without diffusing away, which makes it impossible for BAPTA to intercept Ca^2+^ ions. If so, similar experiments conducted with free Ca^2+^ indicators should report a significantly lower proximal Ca^2+^ signal since the measurement by a diffusible dye would effectively spread the signal over a larger volume. Indeed, this is what we found. As shown in **Fig. 5E**, the apparent proximal Ca^2+^ signal in response to a 10-ms brief pulse measured with Cal520LA-free in the presence of 2 mM BAPTA was 0.9 ± 0.2 (n=9), 4-fold lower than proximal Ca^2+^ signals measured with Cal520LA-RBP (3.6 ± 1, n=20). The latter value is closer to the true measure of the Ca^2+^ concentration in the vicinity of the ribbon base. These findings emphasize that the reliable measurement of ribbon-proximal Ca^2+^ signals in the vicinity of Cav channels greatly benefits from the increased spatial resolution of our nano-physiology approach. We also measured the spatiotemporal properties of local Ca^2+^ signals using Cal520HA-RBP as we have done above for Cal520LA-RBP (**Supplementary Fig. 3**). We found that the ratio between proximal and distal Ca^2+^ signals was similar in 0.2 and 2 mM EGTA (proximal-to-distal ratio ∼1.5, **Supplementary Table 2**) but was enhanced with 10 mM EGTA (proximal-to-distal ratio ∼2.6, **Supplementary Table 2**) and further enhanced with 2 mM BAPTA (proximal-to-distal ratio ∼2.8, **Supplementary Table 2**). Cal520 acts as a buffer that binds Ca^2+^ and carries it away by diffusion. Thus, the higher affinity indicator Cal520HA will bind and shuttle away more Ca^2+^ than Cal520LA. If the Cal520HA indicator is saturated near the Cav channels, it may be underreporting the fast Ca^2+^ transients that occur in those locations. However, Cal520LA and Cal520HA show similar increase in the ribbon proximal-to-distal ratios with increasing concentrations of exogenous buffer.

### Computational models of Ca^2+^ signals along the axis of the RBC synaptic ribbon

Since we could identify the position of the ribbon and we used RBP-fused Ca^2+^ indicator-RBP, we were able to measure and distinguish ribbon-proximal vs. ribbon-distal Ca^2+^ signals that drive the release of UFRP and RRP. However, several factors, including finite spatiotemporal resolution of optical imaging, spatial diffusion, dye saturation, and dye binding kinetics, may limit our ability to achieve optimal resolution. These limitations, however, can be overcome by the use of quantitative models. Thus, we developed a model based on our data with published information on Ca^2+^ diffusion and buffering to estimate more accurately the [Ca^2+^]_I_ gradient along the ribbon at distinct distances from the plasma membrane.

Combining whole-terminal Ca^2+^ current measurements and our estimate for the number of synaptic ribbons per terminal allowed us to infer the magnitude of Ca^2+^ current per single ribbon, which we used in our model to determine the spatiotemporal [Ca^2+^] dynamics near the ribbon. **Fig. 7** shows the results of the simulation of [Ca^2+^] at various distances from the ribbon during and after a depolarizing pulse of 10ms duration in the presence of different concentrations of exogenous and endogenous buffers, replicating the conditions used in our experiments. The geometry of the simulation domain box is shown in **Fig. 6**; it represents the fraction of total synaptic terminal volume per single ribbon (see *Methods*, Supplementary **Table 3** and **Supplementary Fig.4A**.). **Fig. 7** shows the time-dependence of [Ca^2+^] during and after the depolarization pulse at 5 specific locations both near and distant from the ribbon (left-hand panels in each subplot). These five spatial locations are marked by circles of different colors in the right-hand panel of each subplot, which shows [Ca^2+^] at the end of the Ca^2+^ current pulse in pseudo-color in the entire planar cross-section of the simulation domain cutting through the middle of the ribbon, as shown in **Fig. 6**. We assumed a highly simplified Cav channel arrangement into four clusters forming a square with a side length of 80nm. The closest of the five spatial locations (red curves and circles in **Fig. 7**) is X=20 nm away from the base of the ribbon center-line, Z=10 nm above the ribbon, about 45 nm away from the closest Cav channel cluster (see **Fig. 6**). We assumed that this location was within the Ca^2+^ microdomain that triggered the release of the UFRP. Vesicles were not included in the simulation since the total exclusion volume attributed to them represented only a small fraction of the total inter-terminal volume and, therefore, did not significantly impact [Ca^2+^] at the qualitative resolution level we are interested in. A couple of simulations with vesicles included were performed to confirm this statement, requiring much finer spatial resolution and longer computational time. We note also that [Ca^2+^] for short pulse durations considered here is relatively insensitive to our assumptions on the membrane Ca^2+^ extrusion mechanisms, listed in **Supplementary Table 3.**

**Fig. 6.**
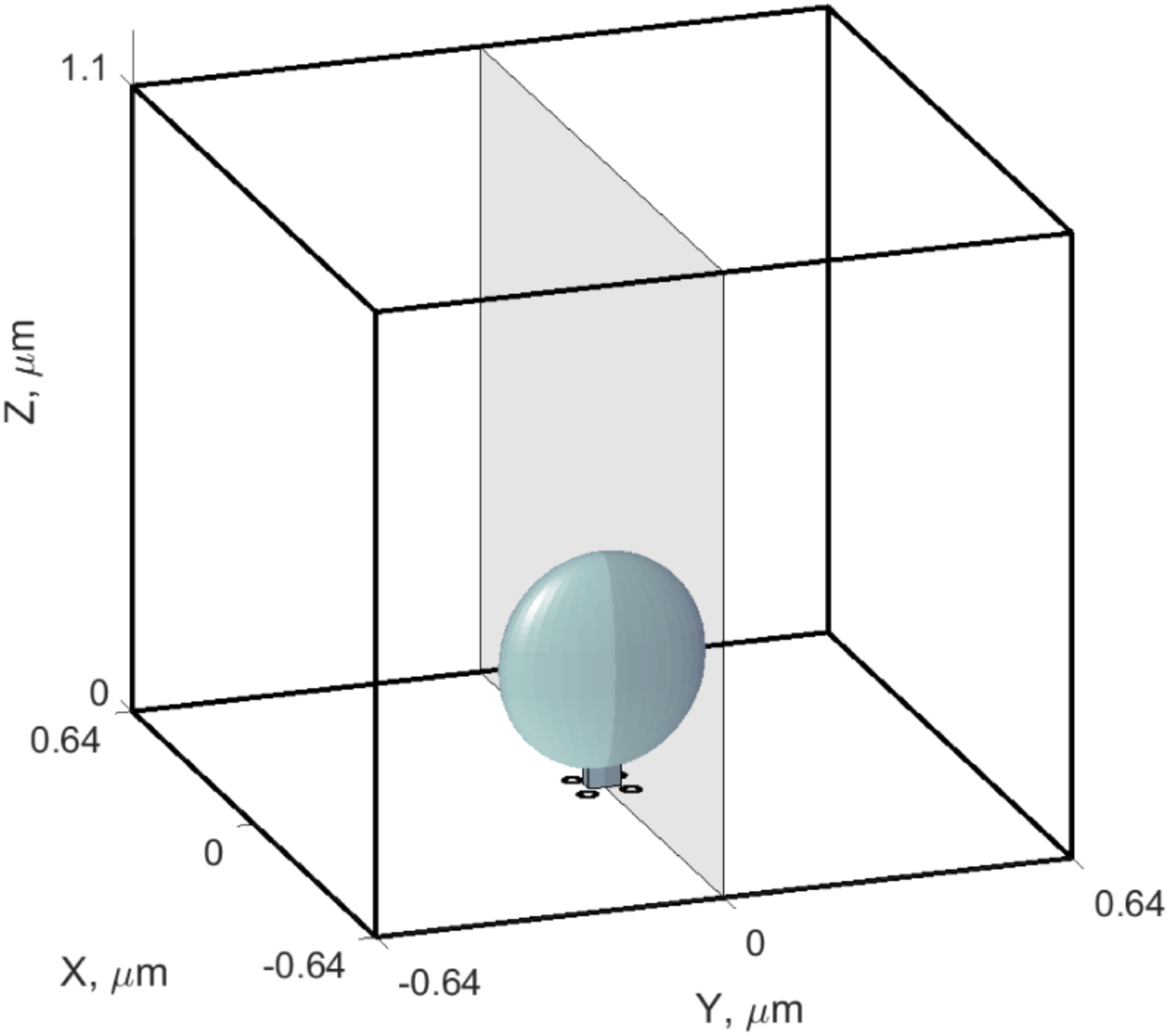
Geometry of the computational model of intra-terminal Ca^2+^ dynamics. Simulation domain is a box with dimensions (1.28 × 1.28 × 1.1) µm^3^, approximating the fraction of synaptic terminal volume per single ribbon. Ca^2+^ ions enter near the base of the ellipsoidal ribbon at four locations marked by black disks representing Ca^2+^ channels or their clusters. The semi-transparent gray coordinate plane Y=0 corresponds to the section used for the pseudo-color 2D [Ca^2+^] plots in **Figure 7**. Ca^2+^ is extruded on all surfaces of this domain, simulating combined clearance by pumps and exchangers on the plasmalemmal as well as internal endoplasmic reticulum and mitochondrial membranes.

**Fig.7.**
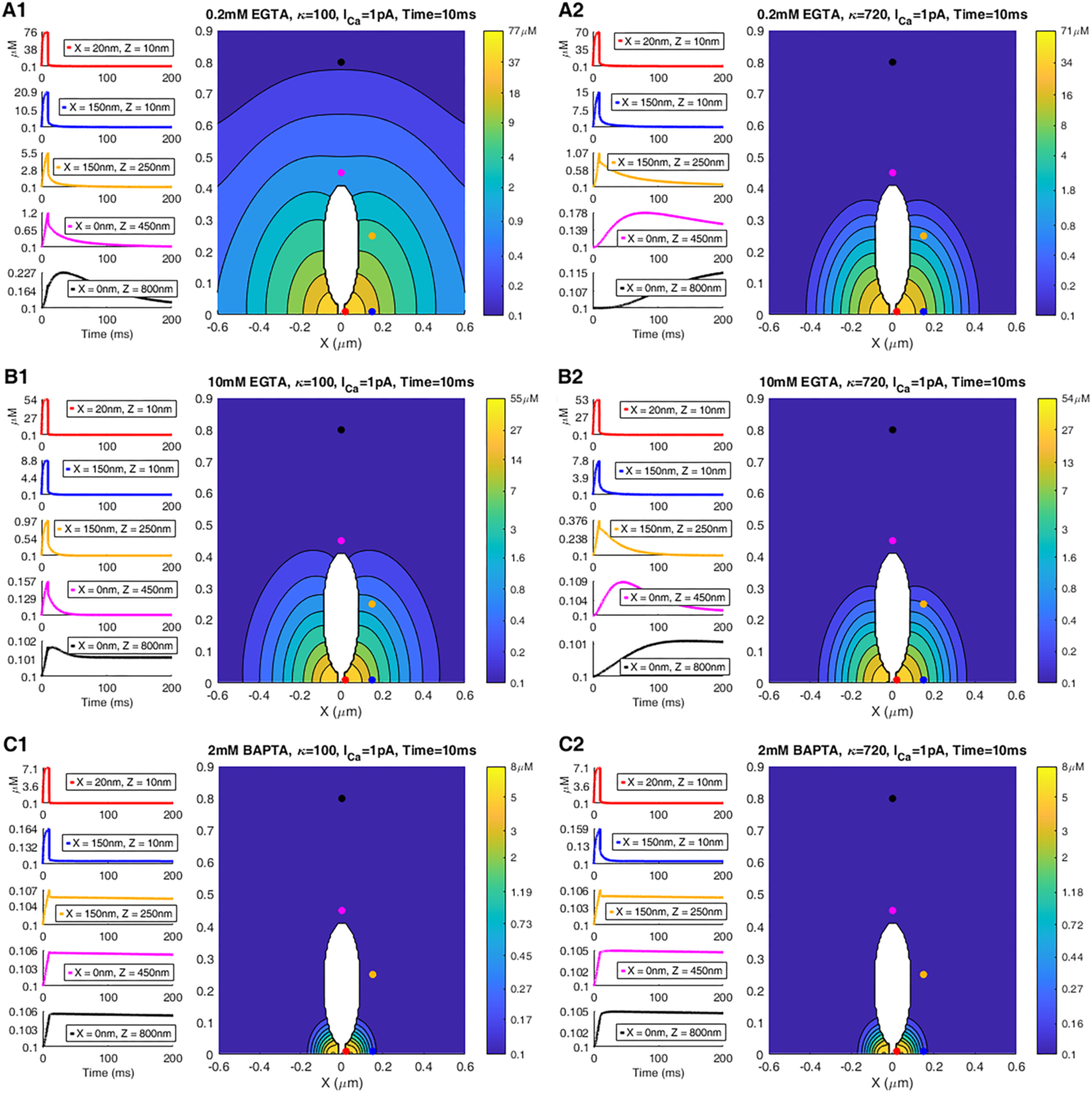
Simulation of the effect of an endogenous immobile buffer of different concentrations on [Ca^2+^] dynamics in response to a 10ms pulse. (**A1-C2**) In each sub-plot, the left-hand panels show the [Ca^2+^] time course in response to a 10 ms constant current pulse (total current of 1 pA) at five select locations marked by colored circles in the right panel. The right-hand panels show a pseudo-color plot of [Ca^2+^] in a 2D section of the 3D simulation volume in **Fig. 6**, at a fixed point in time corresponding to the end of the current pulse. Concentration values for each level curve are indicated in the color bar. Endogenous buffer is immobile, with a concentration of 200 µM in **panels A1-C1** (resting buffering capacity 100 µM), vs. 1.44 mM in **panels A2-C2** (resting buffering capacity 720). Exogenous buffer concentrations were 0.2 mM EGTA (**panels A1, A2**), 10 mM EGTA (**panels B1, B2**), 2 mM BAPTA (**panels C1, C2**).

**Fig. 7A1-B2** reveals that [Ca^2+^] could rise above 50 µM within the microdomain near the base of the ribbon in the presence of 0.2 mM up to 10 mM EGTA. Given the size of the microscope point-spread function, this is in qualitative agreement with the 26 μM estimate of ribbon-proximal [Ca^2+^] that we recorded using the ribeye-bound low-affinity indicator. In addition, this concentration level is reached very soon after the channel opening event due to the rapid formation of the Ca^2+^ microdomain ^76,77^, and therefore, this estimate is expected to hold for shorter pulses as well. However, [Ca^2+^] decayed rapidly with distance, with the rate of decay significantly increasing when EGTA concentration was increased to 10mM (**Fig. 7B1, B2**). Note that a 7-fold change in the capacity of the immobile endogenous Ca^2+^ buffer (cf. **Fig. 7A1 vs. 7A2**) had a relatively modest effect on the [Ca^2+^] level up to about 150 nm distance from the ribbon base. This effect of increasing endogenous buffer concentration was further reduced in 10 mM EGTA (cf. **Fig. 7B1 vs. 7B2**), due to strong competition of endogenous buffer with large concentrations of EGTA. This agrees with the expectation that immobile buffers primarily slow down Ca^2+^ signals but do not affect the rapidly forming quasi-steady-state Ca^2+^ microdomains in the immediate vicinity of the channel ^76–78^. Finally, **Fig. 7C1-C2** shows that 2 mM BAPTA had a much greater effect on localizing the Ca^2+^ signal to the immediate vicinity of the ribbon base.

Since a native unpatched cell may well contain mobile rather than immobile buffers, it was interesting to examine the effect of increasing the concentration of mobile endogenous buffer on Ca^2+^ dynamics in the ribbon vicinity, as compared to the corresponding effect of immobile buffer. **Supplemental Figure 4B-C** shows that a 7-fold increase in mobile buffer concentration reduces the microdomain Ca^2+^ by a factor of 2 and greatly reduces the size of the microdomain. This contrasts with the effect of increasing the concentration of the immobile buffer, which has a much more subtle effect (cf. **Fig. 7A1, A2**). We note that 0.2 mM EGTA was absent in the simulation with mobile buffer shown in **Supplemental Figure 4B-C**; in general, the effect of modest concentrations of EGTA is expected to be negligible due to its slow Ca^2+^ binding speed compared to the endogenous buffer.

### Variability of local Ca^2+^ signals across the RBC synaptic ribbons

Bipolar cells release neurotransmitters primarily from ribbon active zones ^1,6,8,11–13,27–31^. However, the factors that shape the synaptic ribbon microdomains at retinal ribbon synapses have not been examined. We wondered whether all of the 30∼50 synaptic ribbons at an RBC terminal release glutamate in a similar fashion or whether there is some heterogeneity in glutamate release from different ribbon active zones. In particular, we asked what underlying mechanisms could differentiate the Ca^2+^ signals between ribbons of the same RBC terminal and across different RBCs. We first found evidence for such variability in local Ca^2+^ signals at single-ribbon locations of different cells using the freely diffusible high-affinity indicator Cal520HA, observing high variability in the Ca^2+^ transient amplitudes, even for RBCs with similar depolarization-evoked Ca^2+^ current amplitudes (cf. **Supplementary Figs. 5A** vs. **5B**). However, we did not observe such variability when we obtained multiple line scans across the same ribbon (**Supplementary Figs. 5A** vs. **5B**, gray traces). We wondered whether the observed variability between cells could be due to distinct subtypes of zebrafish RBCs ^79^, which might have different subtypes or numbers of Cav channels near the ribbon. Recent work in zebrafish retina identified two distinct RBC subtypes: RBC1 and RBC2 ^79^. The wiring pattern of amacrine cells postsynaptic to RBC1 closely resembles the circuitry of mammalian RBCs, whereas RBC2 forms distinct pathways. Furthermore, RBC1 specifically labels for the known marker of mammalian RBCs, PKC-α ^79^. Due to these similarities, RBC1 in zebrafish is classified as analogous to mammalian RBCs. Moreover, RBC1s have expected morphological characteristics, in particular, the shape and size of soma and the presence of a single synaptic terminal ^80^. Immunolabeling of isolated RBC preparation used in these experiments showed primarily intact RBCs, which are PKC-α-specific RBC1 (**Supplementary Fig. 2B**). For simplicity, and because this study focuses exclusively on RBC1, we will refer to RBC1 as ‘RBC’ throughout this work. Thus, we attribute the variability in the RBC Ca^2+^ transients to the variability between the Ca^2+^ microdomains within the same cell type.

To identify mechanisms that contribute to the heterogeneity in the local Ca^2+^ signals we have reported in this study, we began by asking whether differences in local Ca^2+^ buffering could account for this heterogeneity. To examine the role of Ca^2+^ buffering in the variability of local Ca^2+^ signals across different RBC and different ribbons (**Figs. 8A** and **8D**, respectively), we compared proximal (**Figs. 8B** and **8E**) and distal (**Figs. 8C** and **8F**) Ca^2+^ signals in 10 mM EGTA by averaging the Ca^2+^ recordings from all the ribbons of a given RBC (**Supplementary Figs. 6** and **7**). Here, we first compare proximal (panel B) and distal (panel C) calcium signals across several RBCs, labeled RBC-a through RBC-d. Each RBC contains multiple ribbons, and for each cell, we present the average calcium signals from multiple ribbons using box plots in panels B and C. In these box plots, the horizontal lines represent the average calcium signal for each cell, while the size of the error bars reflects the variability in proximal and distal calcium signals among the ribbons within that RBC. For example, RBC-a had five identifiable ribbons. When the Ca^2+^ signals were restricted to the ribbon sites with 10 mM EGTA in the pipette solution (**Supplementary Tables 1** and **2**), the amplitude of the ribbon-proximal and ribbon-distal Ca^2+^ signals averaged over all ribbons of a given cell exhibited significant variation across different RBCs (**Figs. 8B, 8C**, and **Supplementary Fig. 6**). In panels **D–F (Fig. 8)**, we use RBC-a to illustrate the variability in Ca^2+^ signals across individual ribbons. Specifically, we distinguished proximal and distal Ca^2+^ signals from five ribbons (ribbons 1–5) within RBC-a. We next compared variability in Ca^2+^ signals in the presence of 10 mM EGTA at individual ribbons within the same cell (**Fig. 8D** and **Supplementary Fig. 7**) at proximal (**Fig. 8E**) and distal (**Fig. 8F**) locations. The box plots in panels **E** and **F (Fig. 8)** display the average Ca^2+^ signal (horizontal lines) for each ribbon, based on multiple recordings. For the cell described in **Figs. 8D-8F**, the proximal Ca^2+^ signals were significantly different across all ribbons examined, and there were considerable differences in distal Ca^2+^ signals between ribbons, with the exception of ribbons numbered 2 and 5. Importantly, the lack of or minimal error bars for repeated measurements at the same ribbon indicates that the proximal and distal calcium signals are consistent within a ribbon. These findings emphasize that the observed variability among ribbons and among cells reflects true biological heterogeneity in local calcium domains, rather than experimental noise. The heterogeneity in proximal and distal Ca^2+^ signals at distinct ribbons within the same cell may result from different underlying mechanisms, for example, heterogeneity in Cav expression or subtype and Ca^2+^ handling mechanisms. These findings suggest that exogenous Ca^2+^ buffering has a negligible effect on experimentally observed heterogeneity and variability of the proximal Ca^2+^ signals and that local Ca^2+^ signals at RBC ribbons are dominated by Ca^2+^ in regions close to the ribbon base where Cav channels are located.

**Fig.8.**
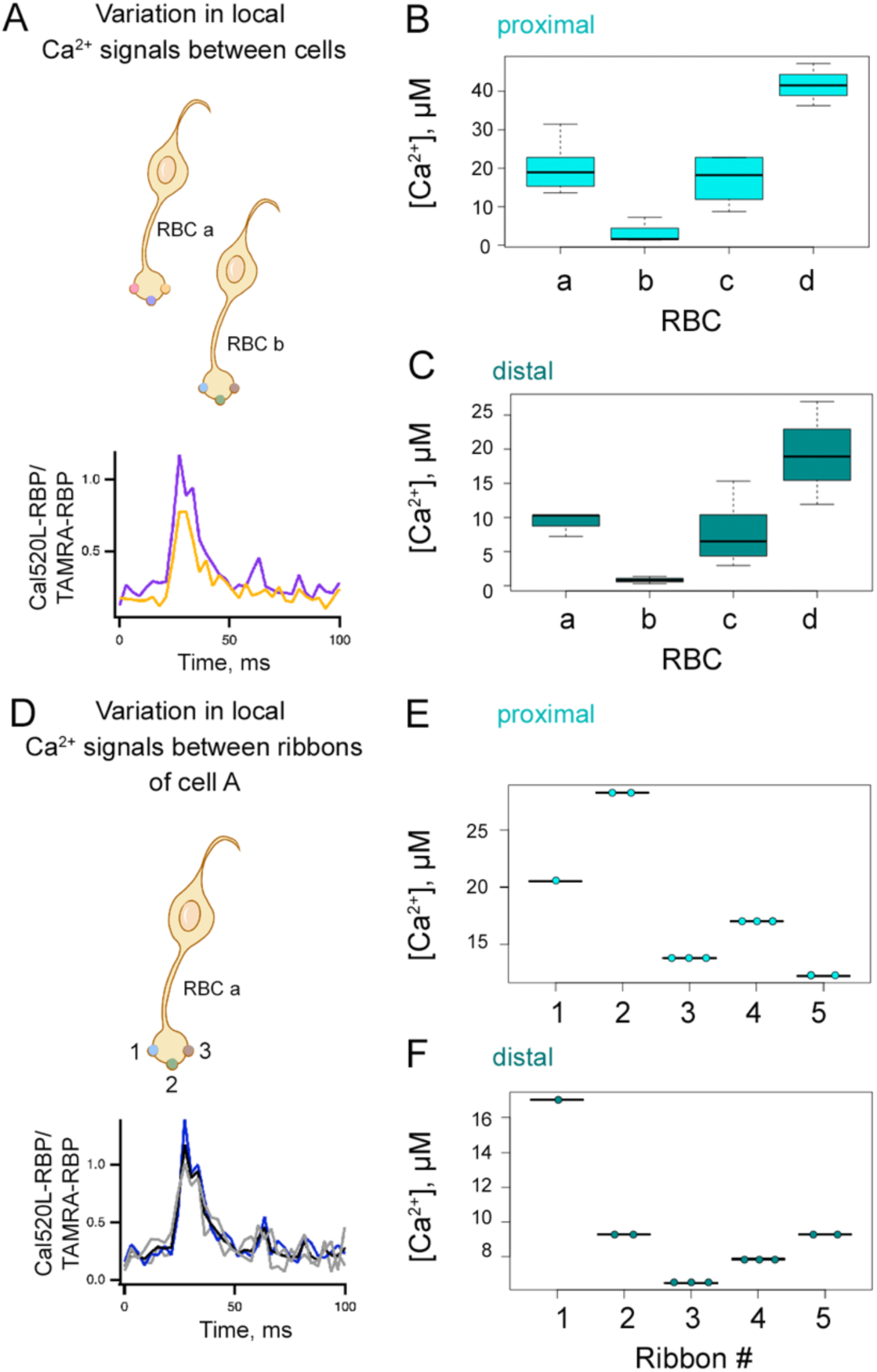
Heterogeneity in the spatiotemporal properties of Ca^2+^ microdomains in RBC terminal. **A. Top panel.** Cartoon of two representative RBCs (cell A and cell B), each containing differently colored ribbons. **Bottom panel.** Ribbon-to-ribbon variability was measured by recording local Ca^2+^ signals near different ribbons (yellow and purple traces) for each RBC. If an RBC had multiple readings for a single ribbon, averages were obtained for comparisons as described in **Supplementary** Fig. 6. **Bottom panel inset:** sample Ca^2+^ currents for the cells from which the Ca^2+^ signal sample traces mentioned above were obtained (purple and yellow traces). Currents were similar across the different cells. Vertical scale = 80 pA, horizontal scale = 5 ms. **B-C.** Variability of Ca^2+^ signals in different ribbons across different cells in the presence of 10 mM EGTA, recorded using Cal520LA-RBP. Proximal Ca^2+^ amplitude values were significantly different between cells A and B (p = 0.022), cells A and D (p = 0.023), and cells B and D (p = 0.005) but similar between A and C (n = 4 cells, 4 fish), as assessed by unpaired t-tests. (**C**) In distal locations, ribbon amplitude values were significantly different between cells A and B (p = 0.031), but similar across all other cell comparisons (n = 4 cells, 4 fish), as assessed by unpaired t-tests. The currents were not significantly different across the different cells, as assessed by unpaired t-tests (mean currents in RBC a: 47.7 ± 2.9 pA, RBC b: 43.7 ± 4.0 pA, RBC c: 43.4 ± 2.0 pA, RBC d: 40.9 ± 2.4 pA; RBC a vs. RBC b: p = 0.45, RBC a vs. RBC c: p = 0.22, RBC a vs. RBC d: p = 0.14, RBC b vs. RBC c: p = 0.93, RBC b vs. RBC d: p = 0.54, RBC c vs. RBC d: p = 0.52). **D. Top panel**. Illustration of a RBC containing three ribbons (numbered 1-3). **Bottom panel.** Ca^2+^ signal measurements from three distinct ribbons (black, gray, and blue traces) were compared to determine the ribbon-to-ribbon variability within each RBC, as described in **Supplementary** Fig. 7. **Bottom panel inset:** sample Ca^2+^ currents for the cells from which the Ca^2+^ signal sample traces mentioned above were obtained (black, gray, and blue traces). Currents were similar across the different cells. Vertical scale = 80 pA, horizontal scale = 5 ms. **E-F.** Box plot illustrating [Ca^2+^] across various ribbons of an individual RBC, which is shown as RBC a. Ribbon variability within individual cells was measured with 10 mM EGTA using Cal520LA-RBP at (**E**) proximal and (**F**) distal locations. (**E**) Proximal Ca^2+^ amplitude values were significantly different among all ribbons (paired t-test, p < 0.001) (n = 5 ribbons, 1 RBCs, 1 fish). (**F**) Distal Ca^2+^ amplitudes were significantly different among all ribbon comparisons (paired t-test, p < 0.001) except for ribbons 2 and 5 (n = 5 ribbons, 1 RBCs, 1 fish). Similar analyses were conducted in two more cells and found similar observations (data not shown). The currents were similar across all ribbons since these were readings from the same cell. Given that some ribbons only have one reading, it is not possible to conduct a paired t-test to statistically compare them; however, the average current ± standard error for the cell shown was 47.7 ± 2.9 pA.

### The ultrastructure of the zebrafish RBC terminal reveals diversity in the size of the synaptic ribbons across the terminal

In hair cells, Ca^2+^ microdomain signaling varies with ribbon size, reflecting larger patches of Cav channels aligned with larger ribbons ^57^. However, the number, size, and shape of ribbon active zones per terminal have not been established in our experimental system, the synaptic terminals of zebrafish RBCs. To provide quantitative measurement of ribbon microdomains, we examined the ultrastructure of zebrafish RBC synaptic ribbons by serial block face scanning electron microscopy (SBF-SEM) and reconstructed three RBCs from serial section electron micrograph (**Fig. 9A**). We analyzed the bipolar cell terminals that are closest to the ganglion cell layer with a morphology similar to mammalian RBCs and zebrafish RBC1s ^79^. Our reconstruction of three RBCs revealed 30-41 ribbons within RBC terminals (**Fig. 9A**), similar to what was previously reported in goldfish giant ON-type mixed RBCs (range 45-60 ^81^).

**Fig. 9.**
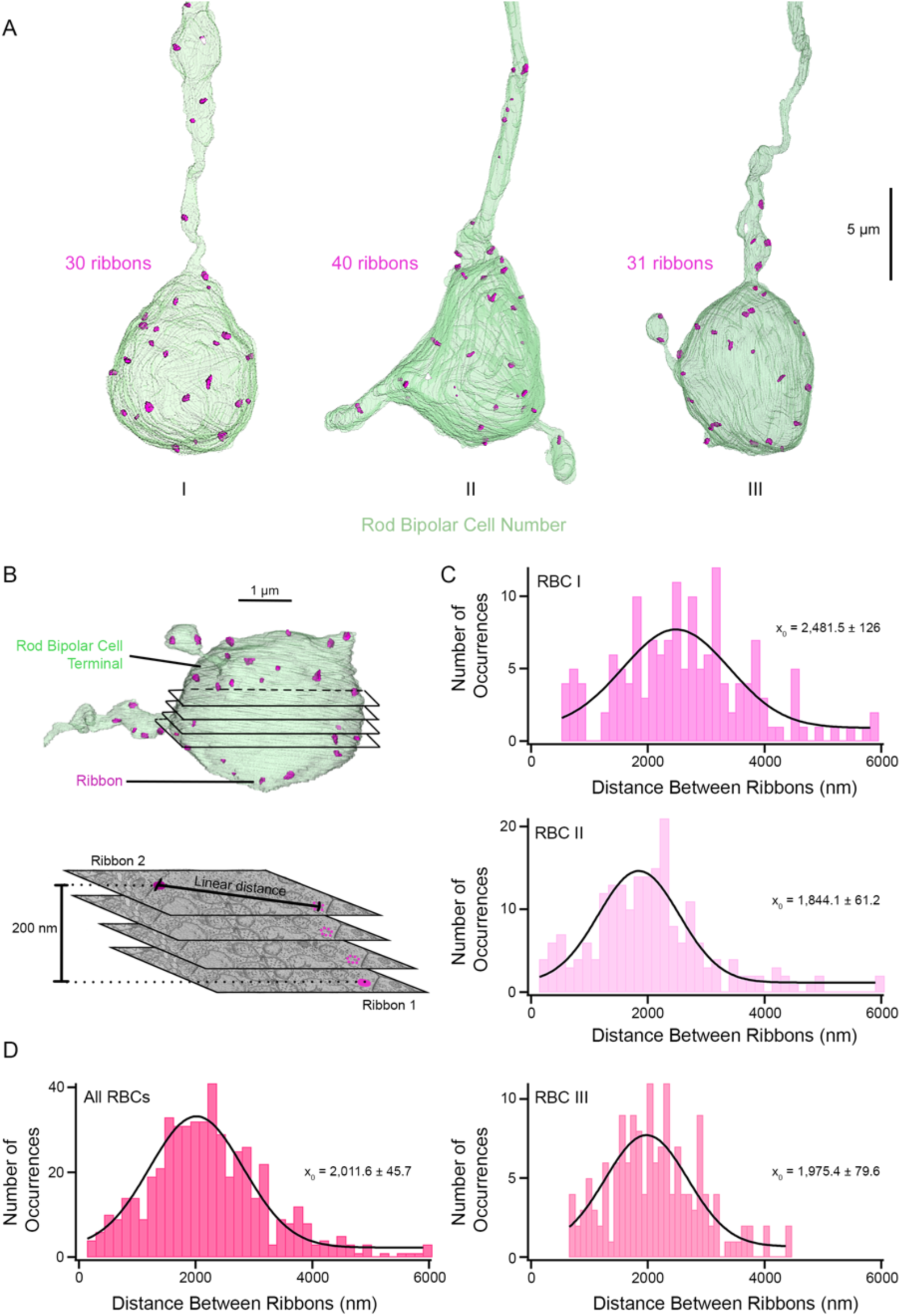
Distribution of measured distance between synaptic ribbons in three zebrafish RBCs. **A.** Reconstruction of three RBC terminals closest to the ganglion cell layer resembling the shape and size of the mammalian RBC1. The ribbons are shown in magenta. Note that the total number of ribbons includes the “floating” ribbons detached from the plasma membrane. RBC II contained 7 floating ribbons. Scale bar, 5 µm. RBCs, rod bipolar cells. **B.** Overview of the distance measurements. **Top.** 3D rendering of a single RBC terminal is shown in light green with ribbons in magenta. Black lines show an example of the different SBF-SEM layers that are cut to obtain individual images. **Bottom.** Four sample SBF-SEM layers are shown with two example ribbons (magenta) located near each other but in different layers. If two ribbons were on the same layer, their linear distance was taken, whereas if they were located in different layers, their linear distance and height difference were used to calculate their actual distance using the Pythagorean Theorem. Each SBF-SEM layer has a thickness of 50 nm. A 3D volume movie of the synaptic terminal of a zebrafish bipolar neuron synaptic ribbon distribution and measurement is provided in Video 1. **C.** Histograms showing the ribbon distance for the three RBCs. x_0_ shows the mean of the distribution. **D.** Histogram of the distances between each ribbon and its five nearest ribbons for all ribbons contained in the three RBCs. x_0_ shows the mean of the distribution. Please note that the y-axis for **B** and **C** have different sizes, given that they represent the number of occurrences of each event.

We examined the distribution of these 30–41 synaptic ribbons within the zebrafish RBC terminals. The distribution of RBC ribbons as estimated by the distance between ribbon sites (**Fig. 9B** and **Video 1**; see *Materials and Methods*) revealed a wide distribution in synaptic ribbon distance to its nearest neighbor across the three reconstructed RBCs (means of 2.48 ± 0.12 µm, 1.84 ± 0.06 µm and 1.98 ± 0.08 µm; **Fig. 9C**). The distance between RBC synaptic ribbons ranged from 141 nm to 6 µm, with a mean of 2 ± 0.05 µm (**Fig. 9D**; across all three reconstructed RBCs). Of the 510 comparisons of zebrafish RBC synaptic ribbons, only 4 ribbon pairs (0.8%) were separated by distances smaller than our confocal microscope resolution limit of 270 nm.

To reveal whether heterogeneity in synaptic Ca^2+^ signals correlates with active zone size, we compared the shape and size of the ribbon active zones across the three reconstructed RBCs. Our results show significant variations in the shape and size of zebrafish RBC synaptic ribbons in RBCs and associated active zones (**Fig. 10**). On average, individual ribbons spanned 2–5 consecutive sections, with some located within the axons or closer to the terminal. The SBF-SEM images and 3D projections of ribbon structures (colored in magenta) revealed considerable variability in shape and size across all dimensions, as illustrated in **Figs. 10A–C**. The number of Cav channels per active zone in hair cells from chicken, frog, and turtle varies with the size of the synaptic ribbon ^82^. We thus measured the area of the ribbon facing the plasma membrane where the Cav channels are known to be located to estimate the number of Cav channels per active zone and to determine whether variation in active zone size might plausibly contribute to the heterogeneity in Ca^2+^ signaling. The EM images and 3D projection of the plasma membrane (yellow) and the area of the ribbon facing the plasma membrane (cyan), representing the active zone, are shown in **Figs. 10A-C**. We observed large variability in the area of the ribbon facing the plasma membrane or active zone within the RBCs, with substantial variability in their average distribution across the three RBCs (**Figs. 10C, cyan, 10E, Video 2** and **Supplementary Fig. 9**), suggesting that the number of Cav channels per RBC active zone could plausibly be heterogeneous.

**Fig. 10.**
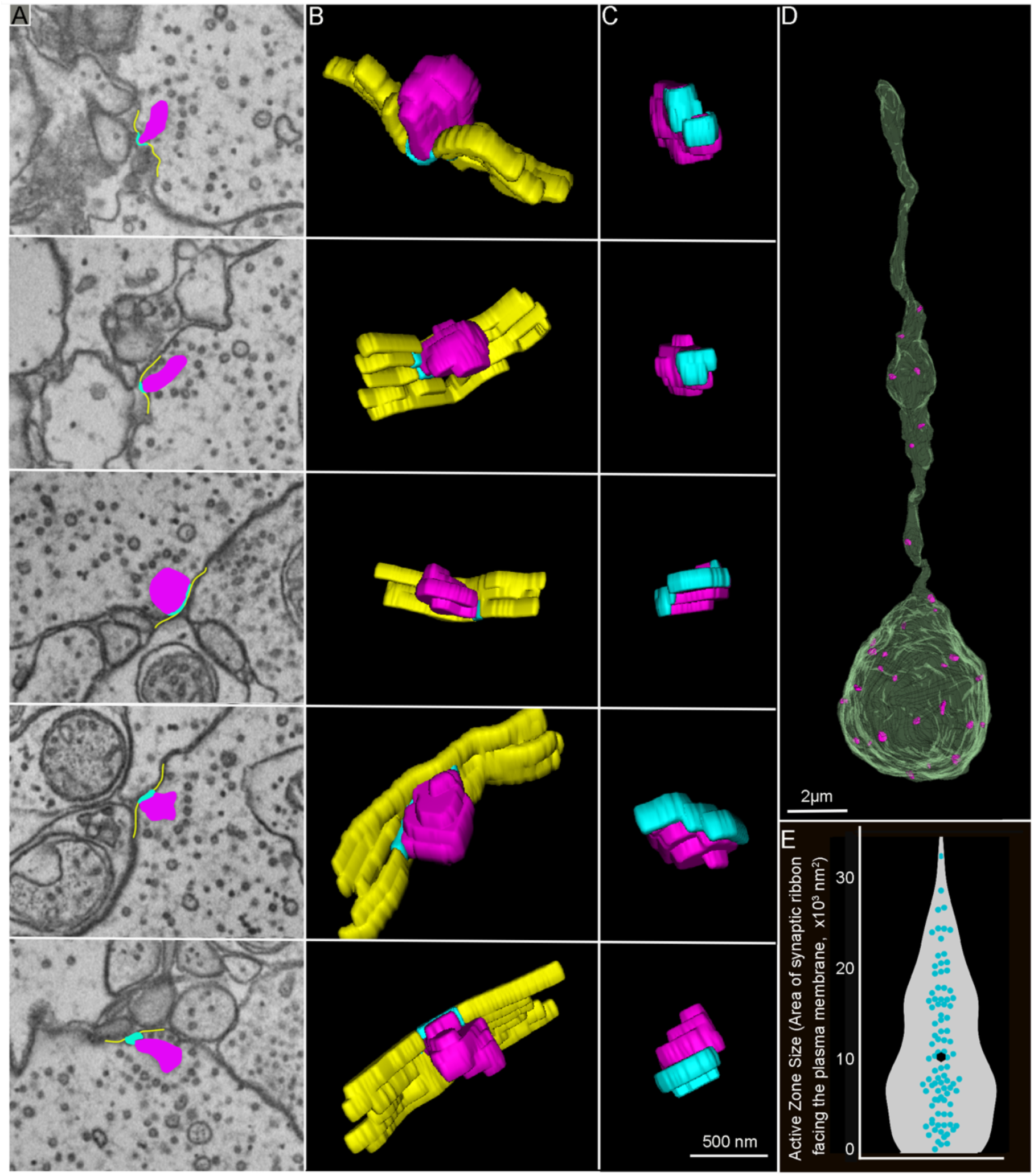
Serial block-face scanning electron microscopy analysis reveals heterogenous RBC ribbon shape, size and area of the ribbon facing the plasma membrane. **A-C.** EM images of zebrafish RBC ribbon structures (**A**) and their respective 3D reconstructions to illustrate different shapes and sizes of synaptic ribbons (**A, B,** and **C**, magenta), plasma membrane (**A** and **B**, yellow), and the area of the ribbon facing the plasma membrane (**A, B,** and **C**, cyan). Each of the 5 rows of images illustrates one ribbon synapse from a zebrafish retinal RBC. A 3D reconstruction of the RBC synaptic terminal and ribbon from SBF-SEM stacks is provided in **Video 2**. **D.** 3D reconstruction of the RBC terminal closest to the ganglion cell layer resembles the shape and the size of the mammalian RBC1 ^2^. The ribbons are colored magenta. **E.** Summary of the active zone size, the area of the ribbon associated with the plasma membrane measured in serial sections across the three RBCs from **Fig 9A**. The individual distribution of the three RBCs active zone sizes is provided in **Supplementary Fig. 8**. The z-size of the SBF-SEM sections is 50 nm, and each ribbon spans 2-5 consecutive sections. The solid cyan circles in the violin plots show individual synaptic ribbon measurements from three RBCs, with the average measurements shown in the solid black circle.

### The size of the active zone and maximal Ca^2+^ influx correlate with the size of the synaptic ribbon

To reveal whether larger ribbons display stronger maximal Ca^2+^ influx, we measured Ca^2+^ signals in response to a series of 200-ms depolarizations. We first imaged TAMRA-RBP and Cal520LA-RBP in sequential scans to obtain ribbon location and resting Ca^2+^ levels. To maximize the capture of Ca^2+^ signals during brief stimuli, we image the Cal520LA-RBP channel, followed by TAMRA-RBP and Cal520LA-RBP sequential scans to confirm the ribbon locations. We found that the maximum amplitude of depolarization-evoked Ca^2+^ signals increased with an increase in ribeye fluorescence (**Fig. 11A & B** and **Video 3;** r = 0.51, N = 121 ribbons, p< 0.001), consistent with findings reported in cochlear inner hair cells ^57,62^. Since ribeye fluorescence correlates with the number of ribeye molecules per ribbon, and, therefore, ribbon size ^32,57,62,83^, our findings suggest that larger ribbons display stronger Ca^2+^ signals as active zone size scales with ribbon size ^62^. Given the diverse shapes of RBC synaptic ribbons (**Fig. 10A & B**, magenta), we measured the longest length and width of ribbons from EM images and compared this with the active zone size. We found that RBC ribbon length and width have a moderate positive correlation (**Fig. 11C**, r = 0.47, N = 102 ribbons, p< 0.001), and both dimensions have a moderate positive correlation with active zone size, albeit the ribbon width (**Fig. 11D**, r = 0.52, N = 102 ribbons, p< 0.001) has a stronger correlation with active zone size than the ribbon length (**Fig. 11D**, r = 0.32, N = 102 ribbons, p< 0.001). These results suggest that heterogeneity in synaptic Ca^2+^ signals correlates with ribbon dimensions and that active zone and the number of Cav channels ^57,62^ scale with ribbon size.

**Fig.11.**
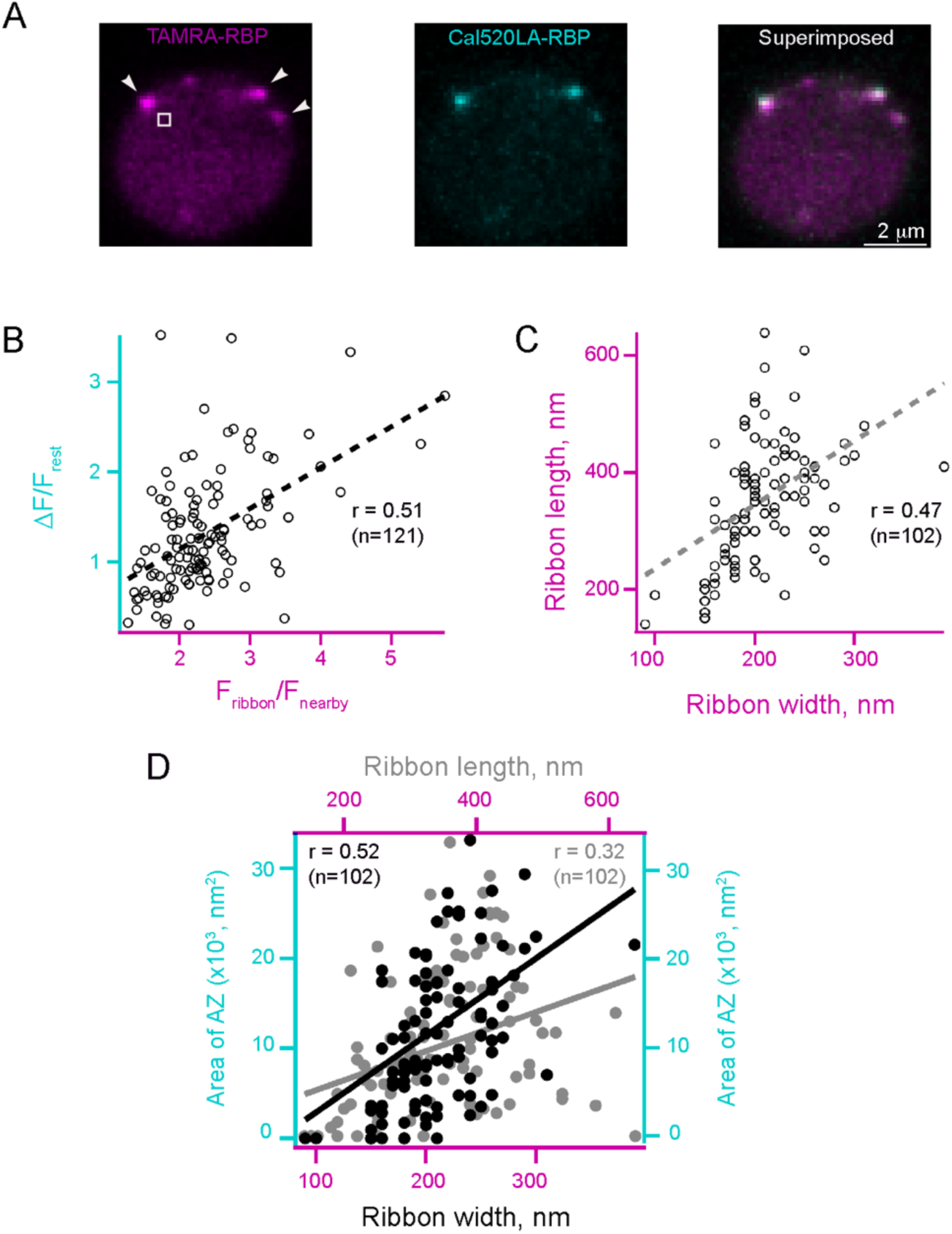
Heterogeneity of Ca^2+^ microdomains in RBC terminals. Larger ribbons have stronger maximal Ca^2+^ influx and larger CAZ. **A.** Images show voltage-clamped RBC filled with a solution containing TAMRA-RBP to label ribbons before depolarization (**left**, magenta) and Cal520LA-RBP to measure the amplitude of Ca^2+^ influx (**middle**, cyan) during depolarization and superimposed of the two (**right**) to compare the size or ribbon vs. maximal Ca^2+^ influx. A time-lapse movie of the synaptic terminal of a zebrafish bipolar neuron during Ca^2+^ influx is provided in Video 3. **B.** Scatter plot of maximal Ca^2+^ influx (F/F_rest_) vs. TAMRA-RBP. Dashed lines are linear regressions, and r is Pearson’s correlation coefficient. **C.** Scatter plot of maximal synaptic ribbon length vs. maximal synaptic ribbon width as estimated from SBF-SEM images. Dashed lines are linear regressions, and r is Pearson’s correlation coefficient. **D.** Scatter plot of area of AZ vs. ribbon width (filled black circles) and of area of AZ vs. ribbon length (filled grey circles). Dashed lines are linear regressions, and r is Pearson’s correlation coefficient. AZ, Active Zone.

We note that a large correlation between the inferred ribbon size and active zone size would be expected if the size heterogeneity is partially due to the differences in the degree of labeling by the two ribbon-binding peptides. Therefore, the observed significant but moderate degree of correlation (r=0.32 or r = 0.52) between the active zone area and the ribbon length or width provides an additional indication of large heterogeneity in the calcium influx between ribbons of similar size, increasing further the heterogeneity in neurotransmitter release by different ribbons.

## Discussion

Ca²⁺ signaling plays a central role in regulating neurotransmitter release throughout the retina, but the spatial and temporal organization of these signals differs markedly between the outer and inner retinal circuits. In the outer retina, photoreceptors rely on graded, sustained calcium influx through L-type Ca²⁺ channels at ribbon synapses to drive tonic glutamate release. These signals are relatively well characterized: the structure– function relationships between ribbon geometry, Ca²⁺ channel distribution, and synaptic vesicle release have been extensively studied, and recent high-resolution studies ^84^ have revealed nanometer-scale alignment between Cav channels and release sites. Postsynaptically, the identity of the glutamate receptor (e.g., mGluR6, KAR, AMPAR) and synaptic cleft geometry further tune the flow of visual information. In contrast, calcium dynamics in the inner retina, especially at bipolar cell terminals, are less well understood. The diversity of bipolar cell types, their complex patterns of ribbon organization, and the presence of both graded and spiking output modes add to this complexity. Moreover, presynaptic Ca²⁺ signals in the inner retina are likely to establish distinct micro- or nanodomain mechanisms depending on ribbon structure, local channel density, and the spatial coupling between Ca²⁺ influx and release sensors. Despite numerous studies that have focused in establishing retinal bipolar cells Ca^2+^ signaling controlling NTR ^9,11,12,48,49,85–87^, there remains significant uncertainty about how presynaptic Ca²⁺ signals are organized across individual ribbon synapses, especially in small or non-classical synapses where standard imaging lacks the resolution to resolve sub-ribbon-scale features. Our study addresses this gap by applying a new immobile Ca²⁺-ribbon indicator, enabling high-resolution tracking of Ca²⁺ domains at the level of individual ribbons. This approach offers a window into nanodomain calcium signaling, allowing us to directly link structural heterogeneity to functional diversity in vesicle release. By capturing the fine-scale dynamics that govern synaptic output in the inner retina, this approach opens the door to a more complete understanding of how visual signals are shaped at their earliest synaptic relays.

### A deep look at the active zone Ca^2+^ microdomain in RBC terminals

In this study, we developed a quantitative nanophysiology approach using ribeye-bound Ca^2+^ indicators and fluorescently labeled RBP, combined with ratiometric dual-color imaging, to measure changes in Ca^2+^ concentration at and along the ribbon axis. By targeting Ca^2+^ indicators to the ribbon combined with fluorescently labeled RBP, we determined the Ca^2+^ concentration along the ribbon at different distances from active zone and found that near the ribbon base the Ca^2+^ concentration could increase to an average of 26 μM upon depolarization. Given the localization of our indicator on the ribbon and minimal sensitivity to exogenous buffers, we believe that our ribbon-proximal signal represents Ca^2+^ concentrations on the ribbon in locations very near release sites, likely some of those that contribute to vesicle resupply. Given that typical resolution for Ca^2+^ imaging—whether using confocal or even STED microscopy—is limited to ∼100–300 nm, the novelty of this study lies in the ability of the new immobile Ca²⁺-ribbon indicator to potentially capture signals on even finer spatial scales, approaching the realm of nanophysiology. Using 2D STED microscopy combined with fluorescence lifetime imaging, Moser’s lab achieved ∼50 nm lateral resolution, enabling precise visualization of Ca²⁺ channel clusters at single active zones in mouse cochlear inner hair cells ^42^. Notably, the [Ca^2+^] _i_ levels obtained from our higher resolution approach using confocal microscopy are similar to those that were reported for measurements in hair cells using Stimulated Emission Depletion (STED) lifetime measurements, emphasizing that our immobile ribbon-bound Ca^2+^ indicator similarly enables nanoscopic (∼tens of nanometers) resolution of localized Ca²⁺ signals—complementing STED-based approaches like Neef et al.’s. ^42^. Previous work on RBCs found slowing of recovery from paired-pulse depression by EGTA, suggesting that Ca^2+^ levels at distal sites on the ribbon might be important for the recruitment of new vesicles following depletion of the RRP and UFRP ^11,30^. Here, we measured directly the effect of EGTA on the spread of Ca^2+^ signals and found that the Ca^2+^ signal at the distal half of the ribbon was two-fold lower than that at the location proximal to the membrane under conditions of low Ca^2+^ buffering. It should be noted, however, that diffraction of light during the optical imaging procedure is likely to underestimate spatiotemporal [Ca^2+^] along the ribbon. Therefore, we used our measurement of the Ca^2+^ current per ribbon to develop a detailed computational model that allowed us to resolve the expected profile of Ca^2+^ microdomains governing UFRP fusion. The model results shown in **Fig. 7** reveal, on a finer spatial scale, the distribution of Ca^2+^ around a ribbon in the presence of various quantities of immobile endogenous buffer and added exogenous buffers. Our finding of high proximal [Ca^2+^] levels on the order of 26 µM recorded using a ribeye-bound low-affinity indicator dye is in qualitative agreement with the simulated high concentration of Ca^2+^ in the microdomain at the base of the ribbon. Simulations also show that, as expected, the peak Ca^2+^ microdomain amplitude is relatively insensitive to the concentration of endogenous immobile buffer and added

EGTA, but its decay with distance is accelerated by increasing the concentrations of exogenous and endogenous buffers. In contrast, the addition of 2 mM BAPTA causes a profound reduction of the Ca^2+^ microdomain amplitude and localizes the signal to a small volume around the ribbon. We also compared the impact of mobile vs immobile endogenous buffers on the Ca^2+^ distribution, showing the much greater effect of mobile buffers on the microdomain amplitude and distance dependence. Although these results are generally expected ^76–78^, the simulation results estimate the [Ca^2+^] levels in the vicinity of the ribbon with greater spatial and temporal resolution. As expected, the ratios of Ca^2+^ concentration values at the base of the ribbon vs. further away from the ribbon are much greater in the model simulation than our experimentally determined proximal-to-distal Ca^2+^ ratio values due to the large size of the microscope PSF, which precludes precise localization of the corresponding Ca^2+^ signals. Further, simulations show the effect of exogenous buffers with greater spatial resolution. However, the simulations fully agree with the overall increase of the proximal-to-distal Ca^2+^ concentration ratios by the application of exogenous Ca^2+^ buffers, especially BAPTA. Despite the technical challenges posed by the limited spatiotemporal resolution of optical imaging, the spatial resolution attained with the nanophysiology approach developed in our study is sufficient to differentiate Ca²⁺ signals associated with distal sites on the ribbon from those localized near the plasma membrane. Conceivably, the nanophysiology approaches established here could generally be applied to other classes of ribbon-containing cells, such as rods, cones and hair cells.

Recent ultrastructural and functional studies have provided new insights into the nanoscale organization of calcium channels at ribbon synapses. In rod photoreceptors, Cav1.4 channels have been shown to localize not only at the base but also along both sides of the synaptic ribbon, in close proximity—within nanometers—to other active zone (AZ) proteins ^84^. This spatial arrangement supports the idea that calcium influx is tightly restricted to subregions of the ribbon, enabling compartmentalized control of synaptic release. These findings are consistent with our own imaging results, which demonstrate localized, submicron Ca²⁺ signals that vary systematically with ribbon geometry. Together, these studies highlight a common theme: that nanodomain Ca²⁺ signaling at ribbon synapses is not only possible but structurally supported by the precise spatial alignment of Ca²⁺ channels and release machinery.

### Heterogeneity of Ca^2+^ microdomains across RBCs terminals

Our quantitative nanophysiology approach provides evidence for the heterogeneity in synaptic Ca^2+^ signals across zebrafish RBCs, suggesting variability in the Ca^2+^ microdomains as has been previously reported for hair cells ^57,62^ and cone photoreceptors. Retinal bipolar neurons contain many ribbon active zones. This ranges from 45-60 in adult goldfish, as quantified by EM ^81^, or 25-45 in RBCs of adult living zebrafish, as estimated from laser scanning confocal microscopy ^88^. Our analyses of ribbon number across zebrafish RBCs through SBF-SEM arrived at an estimate of 31-40. The differences in the number of ribbons as determined through confocal microscopy compared to SBF-SEM could be explained by the resolution limit encountered by the confocal image analyses. Our analyses of zebrafish RBC terminal SBF-SEM images and 3D projections provide the first evidence of diversity in synaptic ribbon shape and sizes across zebrafish RBC terminals. Further, our quantitative measurements of SBF-SEM images reveal that synaptic ribbon width and length correlate with the active zone size, consistent with previous findings in hair cell ribbon synapses ^82,83^ and cone photoreceptor ribbon synapses^58^. Our observation that maximal active zone Ca^2+^ influx correlates with ribeye fluorescence in live imaging is consistent with findings previously reported in hair cells ^62^. Thus, we expect larger active zones, with more Ca^2+^ influx, to have more Cav channel expression, as demonstrated for hair cell ribbon synapses ^62^. Heterogeneity among ribbon active zones has been previously reported in hair cells, where synaptic Ca^2+^ microdomains varied substantially in amplitude and voltage-dependence within individual inner hair cells ^57,62^. Studies in aquatic tiger salamander cone photoreceptors revealed that the amplitude and midpoint activation voltage of Ca²⁺ signals varied across individual ribbons within the same cone. Additionally, local Ca²⁺ signal dynamics at cone photoreceptor ribbons were found to be independently regulated ^58^. However, in comparing our findings with studies of ribbon size heterogeneity in hair cells, we note that we do not observe any systematic variation of ribbon size with larger-scale morphological features like the cell position, as compared to the reported interesting tonotopic variation of ribbon size with position along the cochlea ^57,62^,

The faster, smaller, and more spatially confined Ca^2+^ signals that are insensitive to the application of high concentrations of exogenous Ca^2+^ buffers, referred to here as ribbon proximal Ca^2+^ signals, could be due to Ca^2+^ influx through Cav channel clusters beneath the synaptic ribbon ^25,32,35–48^. However, the variability in Ca^2+^ signals at the distal ribbon, away from the plasma membrane, may result from the spread between two closely spaced microdomains; thus, if proximal Ca^2+^ signals are variable, the distal Ca^2+^ signals are also variable between ribbons. Alternatively, the variability in distal Ca^2+^ signals could be due to additional mechanisms, for example, Ca^2+^ influx from internal stores. Future studies should aim to elucidate the specific mechanisms underlying the observed heterogeneity in distal Ca^2+^ signals. For instance, exploring differences in Ca^2+^ release from intracellular stores or variations in Ca^2+^ sequestration, as reported in goldfish, could reveal key contributing factors ^89^. Although the potential impact of these intracellular Ca^2+^ stores on Ca^2+^ microdomain heterogeneity in hair cells has not been reported,^57^ previous studies in goldfish bipolar cell terminals have documented spatially restricted Ca^2+^ oscillations in voltage-clamped retinal bipolar cells, occurring independently of membrane potential ^26^. To our knowledge, the sources of Ca^2+^ for these oscillations remain unknown. However, previous studies in goldfish bipolar cells suggest that the endoplasmic reticulum^89^ and mitochondria^24^ may act as internal Ca^2+^ stores. Notably, IP₃ receptors have been localized to retinal bipolar cell terminals in some species.^90,91^

In the current study, we investigated possible mechanisms for heterogeneity in proximal Ca^2+^ signals by measuring the ribbon size, particularly the area of the ribbon adjacent to the plasma membrane, where Cav channel clusters are located. Our SBF-SEM shows substantial variability in synaptic ribbon size, shape, and the area of the ribbon facing the plasma membrane where Cav channel clusters are tethered. Since both Ca^2+^ signals and the area of the synaptic ribbon facing the plasma membrane are heterogeneous, we propose that the larger ribbons may anchor more channels, leading to larger Ca^2+^ microdomains. Since local Ca^2+^ signals control kinetically distinct neurotransmitter release components, heterogeneity in local Ca^2+^ signals may alter the rate of vesicle release and allow them to function independently. Indeed, heterogeneity in neurotransmitter release kinetics has been proposed for the observed diversity in excitatory postsynaptic current amplitudes and kinetics reported from paired recordings of goldfish bipolar cells^86^ and ON-type mixed RBCs ^92^. The heterogeneity in the RBC Ca^2+^ microdomains, synaptic ribbon shape, size, and active zone area reported in this study may contribute to regulating the dynamic range of RBC ribbon synapses^28,93,94^ – a hypothesis that needs to be tested in future studies.

## Methods

### Rearing of zebrafish

Male and female zebrafish (*Danio rerio*; 16 ∼20 months) were raised under a 14 h light/10 h dark cycle and housed according to NIH guidelines and the University of Tennessee Health Science Center (UTHSC) Guidelines for Animals in Research. All procedures were approved by the UTHSC Institutional Animal Care and Use Committee (IACUC; protocol # 23-0459).

### Isolation of zebrafish retinal RBCs

Dissociation of RBCs was performed using established procedures ^95^. Briefly, retinas were dissected from zebrafish eyes and incubated in hyaluronidase (1100 units/ml) for 20 minutes. The tissue was washed with a saline solution containing 120 mM NaCl, 2.5 mM KCl, 0.5 mM CaCl_2_, 1 mM MgCl_2_, 10 mM glucose, and 10 mM HEPES, pH = 7.4] before being cut into quadrants. Each quadrant was incubated at room temperature for 25 to 40 min in the same saline solution, to which was added DL-cysteine and papain (20-30 units/ml; Sigma Millipore, St. Louis, MO) and triturated using a fire-polished glass Pasteur pipette. Individual cells were transferred to glass-bottomed dishes, allowed to attach for 30 minutes, and washed with saline solution before being used for experiments.

### Ribeye binding peptides

As a means of localizing the ribbons, custom peptides containing the ribbon binding sequence fused to TAMRA (tetramethylrhodamine; TAMRA: GIDEEKPVDLTAGRRAG) dye were synthesized, purified, and purchased from LifeTein (>95% purity, LifeTein LLC, NJ).

### Ca^2+^ indicators

#### Free Ca^2+^ indicators

The potassium salts of high and low-affinity Ca^2+^ indicators Cal-520^®^ (high affinity, *K_D_* 320 nM) and Cal-520N™ AM (low affinity, *K_D_* 90 µM), referred to as Cal520HA and Cal520LA, respectively were purchased from AAT Bioquest.

#### Direct conjugation of Ca^2+^ indicator to cysteine-containing ribbon-binding peptides

To target Ca^2+^ indicators to the ribbon, custom-made cysteine-containing ribeye binding peptides NH_2_-CIEDEEKPVDLTAGRRAC-COOH were synthesized and purchased from LifeTein to directly fuse with the fluorogenic 520^®^ maleimide (purchased from AAT Bioquest) for high affinity (HA) and low affinity (LA Ca^2+^ indicator dyes). Each peptide at one mM concentration was mixed and incubated with two mM Cal-520^®^ maleimide (20 mM stock solution in DMSO purchased from AAT Bioquest) for one hour at room temperature, then overnight at 4°C. Calibration of Ca^2+^ indicator dyes was performed as described ^25^ and as detailed below. The conjugated Ca²⁺ indicators were stored at -20°C in smaller aliquots, each sufficient for a single day’s experiments.

### Measurement of dissociation constants (K_D_) for Ca^2+^ indicator peptides

The effective *K_D_* (*K*_eff_) was obtained by measuring the fluorescence of Cal520HA-RBP and TAMRA-RBP in buffered Ca^2+^ solutions, determining the ratio between them ^25^, and using the Grynkiewicz equation to determine the Ca^2+^ concentration [Ca^2+^] from this ratio ^59^. However, this was not possible for the low-affinity indicator Cal520LA due to the large Ca^2+^ levels required to calibrate it. Thus, *K*_eff_ for Cal520LA-RBP could be larger than the *K_1/2_* provided by the manufacturer for Cal520LA (*K_D_*, 90 μM), as reported previously with *K_eff_*, measurements of OB-5N in inner hair cells ^42^. Thus, our estimates of local Ca^2+^ concentrations obtained using Cal520LA represent the lower bounds of the underlying true values. As noted in the Results section, the same may be true for the high-affinity Cal520HA despite its accurate *K*_eff_ estimate, due to potential dye saturation effects.

### RBC voltage clamp recording

Whole-cell patch-clamp recordings were made from isolated RBCs, as described previously ^25,33,96^. Briefly, a patch pipette containing pipette solution (120 mM Cs-gluconate, 10 mM tetraethyl-ammonium-Cl, 3 mM MgCl_2_, 0.2 mM *N*-methyl-d-glucamine-EGTA, 2 mM Na_2_ATP, 0.5 mM Na_2_GTP, 20 mM HEPES, pH = 7.4) was placed on the synaptic terminal, as described previously. The patch pipette solution also contained a fluorescently-labeled RBP peptide (TAMRA-RBP) to mark the positions of the ribbons and either 1) free Ca^2+^ indicators Cal520HA-free (**Figs. 1 and 2**) and Cal520LA-free (**Fig. 2, 3 and 5**) to demonstrate our nanophysiological approach or 2) ribeye-bound Ca^2+^ indicators Cal520HA-RBP (**Figs. 3 and Supplementary Figs. 3** and **5**) and Cal520LA-RBP (**Figs. 3, 4, 5, and 8**) to measure local ribbon-associated Ca^2+^ signals. Current responses from the cell membrane were recorded under a voltage clamp with a holding potential (V_H_) of -65 mV that was stepped to 0 mV (t_0_) for 10 milliseconds. These responses were recorded with a patch clamp amplifier running PatchMaster software (version v2×90.4; HEKA Instruments, Inc., Holliston, MA). Membrane capacitance, series conductance, and membrane conductance were measured via the sine DC lock-in extension in PatchMaster and a 1600 Hz sinusoidal stimulus with a peak-to-peak amplitude of 10 mV centered on the holding potential ^97^.

### Acquisition of confocal images

Confocal images were acquired using an Olympus model IX 83 motorized inverted FV3000RS laser-scanning confocal microscopy system (Olympus, Shinjuku, Tokyo, Japan) running FluoView FV31S-SW software (Version 2.3.1.163; Olympus, Center Valley, PA) equipped with a 60 X silicon objective (NA 1.3), all diode laser combiner with five laser lines (405, 488, 515, 561 & 640 nm), a true spectral detection system, a hybrid galvanometer, and a resonant scanning unit. Fluorescently labeled ribeye binding peptide (RBP) ^32^ and Ca^2+^ indicator were delivered to RBC via a whole-cell patch pipette placed directly at the cell terminal. We waited for 30 s after break-in to allow Cal520HA to reach equilibrium with the patch-pipette before obtaining the first fluorescence image. Rapid x-t line scans at the ribbon location were performed to localize synaptic ribbons (**Fig.1Aii**) and to monitor local changes in Ca^2+^ concentration at a single ribbon, as we demonstrated previously, to estimate the Ca^2+^ levels at the plasma membrane and to track a single synaptic vesicle at ribbon locations ^25,26^. The z-projection from a series of confocal optical sections through the synaptic terminal (**Fig. 1Ai**) illustrates ribbon labeling (magenta spots). RBP fluorescence was used to localize a synaptic ribbon and to define a region for placing a scan line perpendicular to the plasma membrane, extending from the extracellular space to the cytoplasmic region beyond the ribbon to monitor changes in the Ca^2+^ concentration along the ribbon axis (**Fig. 1Aii**). The focal plane of the TAMRA-RBP signal was carefully adjusted for sharp focus to avoid potential errors arising from the high curvature near the top of the terminal and the plane of membrane adherence to the glass coverslip at the bottom of the terminal. Sequential dual laser scanning was performed at rates of 1.51 milliseconds per line. Two-color laser scanning methods allowed observation of Ca^2+^ signals (**Fig.1C, cyan**) throughout the full extent of the ribbon in voltage-clamped synaptic terminals, while the ribbon and cell border were imaged with a second fluorescent label. The Exchange of TTL (transistor-transistor logic) pulses between the patch-clamp and imaging computers synchronized the acquisition of electrophysiological and imaging data. The precise timing of imaging relative to voltage-clamp stimuli was established using PatchMaster software to digitize horizontal-scan synced pulses from the imaging computer in parallel with the electrophysiological data (**Fig.1B**). Acquisition parameters, such as pinhole diameter, laser power, PMT gain, scan speed, optical zoom, offset, and step size were kept constant between experiments. Sequential line scans were acquired at 1-2 millisecond/line and 10 μs/pixel with a scan size of 256 × 256 pixels. Bleed-through between the channels was confirmed with both lasers using the imaging parameters we typically use for experiments. To test bleed-through from the RBP channel (TAMRA-RBP) to the Ca^2+^ indicator (Cal-520HA and Cal-520LA), whole-cell recordings from RBC terminals were performed with patch pipette solution that contained TAMRA-RBP or the aforementioned Ca^2+^ indicators, and line-scan images were collected and analyzed using the same procedures used for experimental samples.

### Point spread function

The lateral and axial point spread function (PSF) was obtained as described previously ^25,34,95^. Briefly, an XYZ scan was performed through a single 27 nm bead and the maximum projection in the xy-plane was fit to the Gaussian function using Igor Pro software. We obtained the full width at half maximum (FWHM) values for x and y-width of 268 and 273 nm, respectively, in the lateral (x-y plane) and for y-z width of 561 nm in the axial (y-z axis) resolution.

### Photobleaching

We minimized photobleaching and phototoxicity during live-cell scanning by using fast scan speed (10 microsec/pixel), low laser intensity (0.01–0.06% of maximum), and low pixel density (frame size, 256 × 256 pixels). We estimated photobleaching using x-t line scans of Cal520HA or Cal520LA and TAMRA-RBP in the absence of stimulation, with the same imaging parameters used for experimental samples.

### Data analysis

Quantitative FluoView x-t and x-y scans were analyzed initially with ImageJ software (imagej.nih.gov) and subsequently with Igor Pro software (Wavemetrics, Portland OR) for curve fitting and production of the figures. Data from PatchMaster software were initially exported to Microsoft Excel (Version 16.81) for normalizing and averaging and exported from MS Excel to Igor Pro (Version 9.05) for curve fitting and production of the l figures.

### Analysis of x-t scan data

#### X-axis profile

To determine the Ca^2+^ signals along the ribbon axis, we spatially averaged the x-axis profile intensity of RBP (i.e., a horizontal row of pixels, see **Fig.1C, *top***) to determine the position of the center of the ribbon and estimate the location of the plasma membrane. The parameter *x_0_* is the peak of the Gaussian fit, giving the x-position of the center of the ribbon. The x-axis profile is also used to obtain the spatial profile of Ca^2+^ signals before, during, and after with respect to the ribbon profile. To identify the Ca^2+^ signals specific to the ribbon location, we fit x-axis intensity profiles with the equation *f*(*x*) = *s*(*x*) + *g*(*x*), where *s*(*x*) is a sigmoid function that describes the transition from intracellular to extracellular background fluorescence at the edge of the cell, given by *s*(*x*) = *b* − *c* / (1 − *exp*((*x*_1/2_ − *x*)/*d*)), and *g*(*x*) is a Gaussian function that represents the fluorescence of RBP, given by *g*(*x*) = *a exp*(−(*x* − *x*0)^2^/*w*^2^), as described ^33^. The parameters *x*_1/2_ and *x*_0_ were taken as the x-axis positions of the plasma membrane and the fluorescence emitter, respectively. While parameter b is intracellular background fluorescence, *c* is extracellular background fluorescence, *d* is the slope of the sigmoid, *a* is the peak amplitude of emitter fluorescence, and *w* is √2 times the standard deviation of the Gaussian function, in practice, the latter parameters were highly constrained by the data or by the measured PSF, leaving only *x*_1/2/_ and *x*_0_ as free parameters in the fitting. **Fig.1D**, demonstrates that the peak of the Ca^2+^ signals (*x*_$_, cyan; **Fig.1D**) during the stimuli is proximal to the ribbon center (x_0_, magenta; **Fig.1D**) towards the plasma membrane (*x*_!/#_, magenta; **Fig.1D**), as expected for Ca^2+^ influx originating from Ca^2+^ channels localized in the plasma membrane ^50–52^.

#### T-axis profile

The temporal profiles of the Cal520, Cal520-RBP, and TAMRA-RBP signals were determined by analyzing the time-axis profile of the x-t line scan to obtain the kinetics of the Ca^2+^ transient with respect to the ribbon. We determined the baseline kinetics by averaging the fluorescence obtained immediately before depolarization. The timing of depolarization and the amplitude of the Ca^2+^ current was obtained from the PatchMaster software. The rising phase of the Ca^2+^ transient was fit with the sigmoid function, the peak of which is referred to as the peak amplitude Ca^2+^ transient.

We used this baseline profile of ribbon-proximal Ca^2+^ signals to distinguish signals proximal or distal to the ribbon despite the distance between these two signals being within the PSF. For example, the ribbon-proximal signals were obtained by averaging 5 pixels of the temporal profile of the Ca^2+^ signals (obtained with Cal520 or Cal520-RBP) between the x_1/2_ and x_0_ values obtained in the x-axis profile of TAMRA-RBP. The distal profile was obtained similarly but was 5-pixels after x_0_ towards the cytoplasm.

#### Quantifying the kinetics of the Ca^2+^ transients

The decay phase of the fluorescence transients in Figs. 2 and 3 were fit using the standard bi-exponential sum with time constants *τ_fast_* and *τ_slow_*:

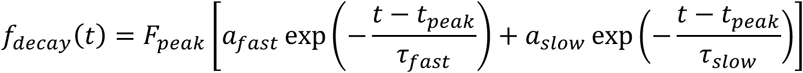

Where *F_peak_* and *t_peak_* are the peak fluorescence and the time at which this peak has been reached, and *a_fast_* and *a_slow_* are the relative proportions of the two decay components, specified as percentages and satisfying *a_fast_* + *a_slow_* 100%. The decay parameters were found using a global differential evolution optimization algorithm ^98^, repeating the optimization 400 times starting with different initial parameters values, to ensure true global minimum of the fit error is reached. We note however that bi-exponential data fitting is known to be an ill-conditioned procedure, highly sensitive to noise and other recording characteristics such as the duration of the measurement. Therefore, much higher signal-to-noise ratio would be required for reliable uncertainty quantification of the corresponding kinetic fit parameters, so we only fit the trial-averages data for fluorescence traces in Figs. 2 and 3 as a rough description of temporal decay of the signal; no precise quantitave conclusions should be drawn from the decay time values. Still, time-course fit of the averaged traces may provide a qualitative-level comparison of stimulus-evoked fluorescence transients recorded with different dyes at different locations with respect to the ribbon.

### Analysis of x-y scan data

We analyzed the rate of loading the TAMRA-RBP and Ca^2+^ indicator into the RBC terminal with ImageJ software by placing a square region of interest (ROI; 5 x 5 μm) and using a Plot z-axis profile function to obtain spatially averaged TAMRA-RBP and Ca^2+^ indicator fluorescence as a function of time. The rising phase of TAMRA-RBP and Ca^2+^ indicator fluorescence was obtained by fitting the rising to the peak to the single exponential function using the curve fitting function in Igor Pro 9 software. The rate of the exponential function is defined as the rate of fluorescence loading to the terminal.

For analysis comparing the fluorescence intensity of ribbons and Ca^2+^ influx elicited by 100-ms stimuli (**Fig. 11A & B**), 10 images of both TAMRA-RBP and Cal520LA-RBP were collected in sequential, followed by imaged Cal520LA-RBP only during 200-ms depolarization, and the sequence ended with 10 images of both TAMRA-RBP and Cal520LA-RBP in sequential to confirm the ribbon locations. Two color imaging were obtained sequentially with a frame interval of 407 ms, and Cal520LA-RBP only during 100ms-stimuli was 205 ms. Images before and after were then averaged to obtain the ribbon location. Synaptic ribbon fluorescence was quantified as the ratio of TAMRA fluorescence to the fluorescence of the nearby RBC cytoplasm, measured approximately eight to nine pixels away (F_ribbon_/F_near by_), measuring the pixel with the highest intensity. The change in Cal520LA-RBP fluorescence, used as a proxy for active zone Ca^2+^ influx, was estimated as ΔF/F_rest_, where F_rest_ is the fluorescence intensity before depolarization at −65 mV, and ΔF is the difference when depolarized to 0 mV. Ca^2+^ indicator intensity was calculated as the average fluorescence of nine pixels centered on the pixel with the greatest fluorescence increase.

### Experimental design and statistical methods

We did not use any statistical methods to determine the sample size prior to experiments. Mean value variances were reported as ± the standard error of the mean (SEM). The statistical significance of differences in average amplitudes of Ca^2+^ current, capacitance, synaptic ribbon size and number, and Ca^2+^ transients were assessed using unpaired, two-tailed t-tests with unequal variance using R Studio (Version 2023.09.0+463) and Igor Pro software.

#### Amplitude analysis for Figure 8

Amplitude data were processed and analyzed using R Studio software (Version 2023.09.0+463). The data were divided into two groups, that of individual cells and of multiple cells, and analyzed as follows. Multiple individual cells from each condition (proximal 10 mM EGTA Cal520LA-RBP and distal 10 mM EGTA Cal520LA-RBP) were analyzed to compare the variability between the Ca^2+^ amplitude measurements of the ribbons from each cell. Having verified the normality of the data to ensure the validity of the statistical test, we used one-way ANOVA and Tukey’s honest significant difference test to assess the specific statistical differences between ribbons. To compare the variability between the amplitude measurements of different cells, we compared readings for various cells. If there were multiple measurements for a single ribbon in a specific cell, those readings were averaged. In this case, the normality of the data was again verified and Welch’s ANOVA was employed to assess the between-group differences as it does not assume homogeneity of variance, with the Games-Howell post-hoc test being used to understand the specific differences between cells.

### Computational modeling of Ca^2+^ dynamics

Spatio-temporal Ca^2+^ dynamics is simulated in a volume shown in **Fig. 9**, which is a 3D box of dimensions (1.28 x 1.28 x 1.1) µm^3^. Assuming an approximately spherical bouton of diameter 5µm and an average of 36 ribbons per terminal, one obtains synaptic volume per ribbon of 1.81 µm^3^, which matches the volume of our box domain. Ribbon serves as an obstacle to Ca^2+^ and buffer diffusion, and has an ellipsoidal shape with semi-axis of 190nm along the Y- and Z-axes (for a total height of 380nm), and a semi-axis of 70 nm along the X-axis. It is attached to the plasma membrane by a thin ridge (arciform density) of dimensions 60×30×30 nm. Ca^2+^ ions enter through 4 channels (or clusters of channels) at the base of the ribbon indicated by black disks in **Fig. 6**, forming a square with a side length of 80 nm; this highly simplified channel arrangement is sufficient to capture the level of detail in spatial [Ca^2+^] distribution that we seek to resolve. Each of the four clusters admits a quarter of the total Ca^2+^ current. Ca^2+^ current values and current pulse durations are listed in **Fig. 7**. Although the actual distribution of Cav channels is more heterogeneous, channels close to the ribbon play a greater role in neurotransmitter release ^2^.

The simulation includes a single dominant Ca^2+^ buffer with molecules possessing a single Ca^2+^ ion binding site. We assume that only immobile buffer is present in a patch-clamped cell, since any mobile buffer would diffuse into recording pipette. Simulations with mobile buffer in an un-patched cell are also provided (**Supplementary Fig. 4B&C**). Buffer parameter values were set to values reported by ^7^, namely dissociation constant of 2 µM, and total concentration of 1.44 mM (total buffering capacity at rest of 720). We also examine the impact of the assumption on buffer concentration by repeating the simulation with a lower buffering capacity of 100 (total buffer concentration of 200 µM). The Ca^2+^ ion diffusivity is set to 0.22 µm^2^/ms ^99^.

To minimize the number of undetermined parameters, we assumed the same parameters for the Ca^2+^ clearance on all surfaces, with one high-capacity but lower affinity process simulating Na^+^/Ca^2+^ exchangers, and one lower capacity but higher affinity process simulating Ca^2+^ extrusion by ATPase PMCA and SERCA pumps. We used the Hill coefficient of *m*=2 for the Ca^2+^ sensitivity of the SERCA pumps on all surfaces except the bottom surface ^100^, whereas Hill coefficient of *m*=1 corresponds to PMCA pumps on the bottom surface ^101^. Thus, the flux of [Ca^2+^] across the surface (ions extruded per unit area per unit time) is described by

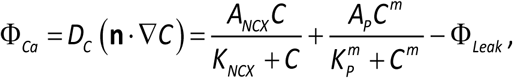

where *C* is the Ca^2+^ concentration, *D_C_* is the Ca^2+^ diffusivity, and ***n*** ⋅ **∇*C*** is the gradient of [Ca^2+^] normal to the boundary. The constant leak term Φ_Leak_ ensures that the flux is zero at the resting value [Ca^2+^]_rest_ = *C*_0_ = 0.1 µM. The Ca^2+^ clearance rate and affinity parameters for the pumps and exchangers are listed in **Supplementary Table 3**. We note however that the density of Na^+^-Ca^2+^ exchangers and PMCA/SERCA pumps are rough estimates and have a significant effect on results only for longer stimulation durations of over 100-200 ms. Reaction-diffusion equations for [Ca^2+^] and buffer are solved numerically using Calcium Calculator software version 6.10.7 ^102,103^.

### Serial block-face scanning electron microscopy (SBF-SEM)

The eyecups of adult (16 ∼20-month-old) zebrafish were dissected in bicarbonate-buffered Ames’ medium and, after removing the lens, the eyecups were fixed in 4% glutaraldehyde in 0.1 M sodium-cacodylate buffer at RT for one hour, followed by overnight fixation at 4°C. Thereafter, samples were treated to generate blocks for serial block face scanning electron microscopy (SBF-SEM) according to the published protocol ^104^, in this case using Zeiss 3-View SBF-SEM. Multiple 35 µm montages were acquired at a 5 nm X-Y resolution and a 50nm Z resolution across the entire retina from the outer (photoreceptor) layer to the inner (ganglion) cell layer. Images were inspected visually and structures of interest were traced using the TrakEM plug-in in ImageJ software (NIH/Fiji), as described below.

#### Analysis of serial block-face scanning electron microscopy images

SBF-SEM image stacks of the zebrafish retina were imported, aligned, visualized, and analyzed using the TrakEM plugin (version 1.5h) in ImageJ software (NIH). RBC1 were identified based on their characteristic morphology and 4-7 μm terminal size, as previously described ^80^. RBCs, synaptic ribbons, pre-synaptic membranes, and the area of the ribbon touching the plasma membrane were traced, painted using the TrakEM brush features, and rendered in 3D for structural visualization. The area of the synaptic ribbon touching the RBC membrane was quantified using the measuring tools in ImageJ and images were prepared using Adobe Photoshop software (version 25.9). The distance between each ribbon from three different RBCs and its nearest five ribbons (**Fig. 9B** and **Video 1**) was measured as follows: Ribbons on the same plane had their linear distance measured, whereas the distance between ribbons that were close to each other but in different layers was calculated using the Pythagorean Theorem.

## Acknowledgements

This research was funded by the National Institutes of Health (NIH) National Eye Institute (NEI), award numbers R01EY030863, 3R01EY030863-02S1, and 3R01EY030863-03S1, and a UTHSC College of Medicine Faculty Research Growth Award (to TV), R01 EY032396, P30EY026878, R01NS122388 (DZ), and a University of Wisconsin-Madison (UW2020) WARF discovery award (to MH) which funded the acquisition of the 3D view serial block-face electron microscope. We would like to acknowledge the UW Madison School of Medicine electron microscopy facility for processing and imaging samples and thank Dr. Alex Dopico, the Van Vleet Chair of Excellence and Professor in the Department of Pharmacology, Addiction Science, and Toxicology, UTHSC, for critical reading, and Dr. Kyle Johnson Moore in the UTHSC Office of Research for editing the manuscript.

## Author Contributions

Conceptualization, D.Z. and T.V.; Methodology and Investigation, N.R., A.S.; Data analysis, N.R., A.S., J.B., M.H., T.V.; Computational modeling, V.M.; Visualization, J.B., M.H., V.M., T.V.; Writing - Original Draft, N.R., T.V.; Writing-Review & Editing, all authors; Funding, M.H., D.Z., T.V.; Supervision and Project administration, T.V.

---

**Video 1.** 3D volume movie of the synaptic terminal of a zebrafish rod bipolar cell terminal showing the distribution of included synaptic ribbons and measurements.

RBC terminal is shown in light green with ribbons in magenta. Illustrations show how the measurements were obtained for ribbons on the same vs. different layers.

**Video 2.** A 3D reconstruction of the RBC synaptic terminal and ribbon from SBF-SEM stacks is provided in Video 2.

**A.** 3D reconstruction of the RBC terminal (green) with included synaptic ribbons (colored magenta).

**B.** 3D reconstruction of the RBC synaptic ribbons shows different shapes and sizes of ribbon (magenta) and the AZ, the area of the ribbon facing the plasma membrane (cyan). AZ, active zone.

**Video 3.** A time-lapse movie of the synaptic terminal of a zebrafish bipolar neuron during Ca^2+^ influx is provided in Video 3.

Left: RBC terminal labeled with TAMRA-RBP shows the locations of synaptic ribbons (magenta spots). Middle: RBC terminal filled with Cal520LA-RBP shows the Ca^2+^ influx in 0.02 s. looped two times to see Ca^2+^ influx in 0.07 s.

Right: Superimposed TAMRA-RBP and Cal520LA-RBP shows the Ca^2+^ influx as spot-like maxima near the membrane during depolarization. TAMRA-RBP 10 images before depolarization, and stacks of Cal520LA-RBP 5 images before depolarization and 5 images during depolarization were compiled to demonstrate the locations and magnitude of the Ca^2+^ influx.

The interval between frames is 407 ms, Total duration of the movie 0.9 s. Each frame is an individual, unaveraged image.

