## Supplemental Figures for "Nanophysiology Approach Reveals Diversity in Calcium Microdomains across Zebrafish Retinal Bipolar Ribbon Synapses"

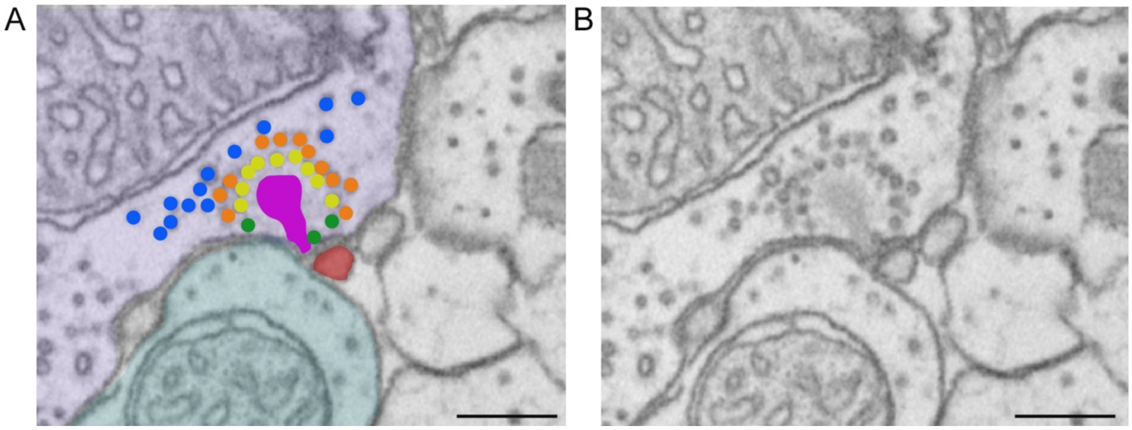


**Supplementary Fig. 1**. **Ultrastructure of a zebrafish RBC showing distinct synaptic vesicle pools.**

**A.** A single section from serial block-face scanning electron microscopy (SBF-SEM) shows RBCs (purple-shaded) and postsynaptic neurons (cyan and brown-shaded) in the SEM volume. Synaptic vesicles in RBC are distributed among at least four distinct pool types based on their fusion kinetics, the average proximity of vesicles to Cav, and the state of vesicle preparedness for $Ca^{2+}$-triggered fusion. All of the vesicles in the ribbon (magenta-shaded) readily releasable pools (RRP) are molecularly prepared for fusion but differ in their anatomical distance to Cav, leading to a clear kinetic distinction between the first and second phases of exocytosis triggered by strongly activating the $Ca^{2+}$ current. The RRP vesicles docked at the base of the synaptic ribbon are defined as the ultrafast releasable pool (UFRP; **panel B**, green vesicles) [1], which are distinct from the remaining RRPs at the ribbon that are distal to the plasma membrane (PM; **B**; yellow vesicles). These anatomically distinct pools contribute to the rapid first phase and slower second phase of neurotransmitter release, respectively [1-5]. The cytoplasmic pool that replenishes the ribbon pools is defined as the recycling pool (RP, orange vesicles), while those that do not participate in neurotransmitter release are members of a reserve pool (Res. P, blue vesicles). Scale bars: 500 nm (**panels A** and **B**).

**B.** The same single SBF-SEM section from **A,** but without shading.

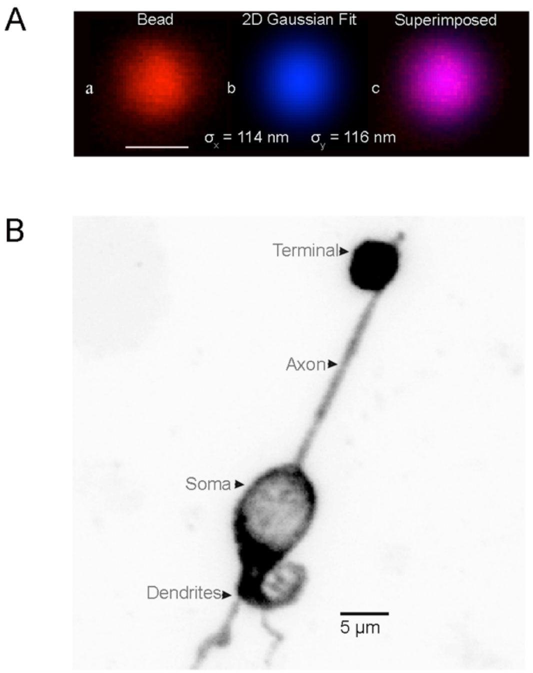


**Supplementary Fig. 2**. **$Ca^{2+}$ indicator fluorescence imaging in the RBC terminal.**

**A. Measurement of the point-spread function (PSF) of the microscope**
**a.** Image of a single 27-nm bead. Scale bar: 250 nm
**b.** Two-dimensional Gaussian fitted to **panel A** using Igor Pro software. The indicated standard deviations (SD) (114 and 116 nm) correspond to a full width at half maximum (FWHM) of 268 and 273 nm in the x and y-axis, respectively.
**c.** Superposition of fit (blue) with bead image (red).

**B.** RBC isolated from zebrafish retina after papain digestion and immunostained with anti-PKCα antibodies shows its characteristic flask-shaped cell body and large terminals.

| | Proximal ($\Delta F/F_{rest}$) | Distal ($\Delta F/F_{rest}$) |
|---|---|---|
| 0.2 mM EGTA | 5.5 ± 0.9 (N=30) | 3.3 ± 0.8 (N=30) |
| 2 mM EGTA | 6.6± 0.8 (N=21) | 4 ± 0.8 (N=21) |
| 10 mM EGTA | 3.5 ± 0.4 (N=43) | 1.8 ± 0.2 (N=43) |
| 2 mM BAPTA | 3.6 ± 1 (N=20) | 0.8 ± 0.2 (N=20) |

**Supplementary Table 1**. **Effect of exogenous $Ca^{2+}$ chelators alter $Ca^{2+}$ signals gradient along synaptic ribbon measured with Cal520LA-RBP.** There were significant differences between proximal vs distal measured as $\Delta F/F_{rest}$, in all conditions as found through paired-sample t-test analysis performed on RStudio. Differences were smaller between proximal vs distal Ca2+ signals in 0.2 mM, 2 mM, and 10 mM EGTA conditions, but more prominent with 2 mM BAPTA (0.2 mM EGTA: proximal vs distal: p = 0.0027, 2 mM EGTA: proximal vs distal: p = 0.034, 10 mM EGTA: proximal vs distal p = 0.00013, 2 mM BAPTA: proximal vs. distal: p = 0.0073, n = 22).

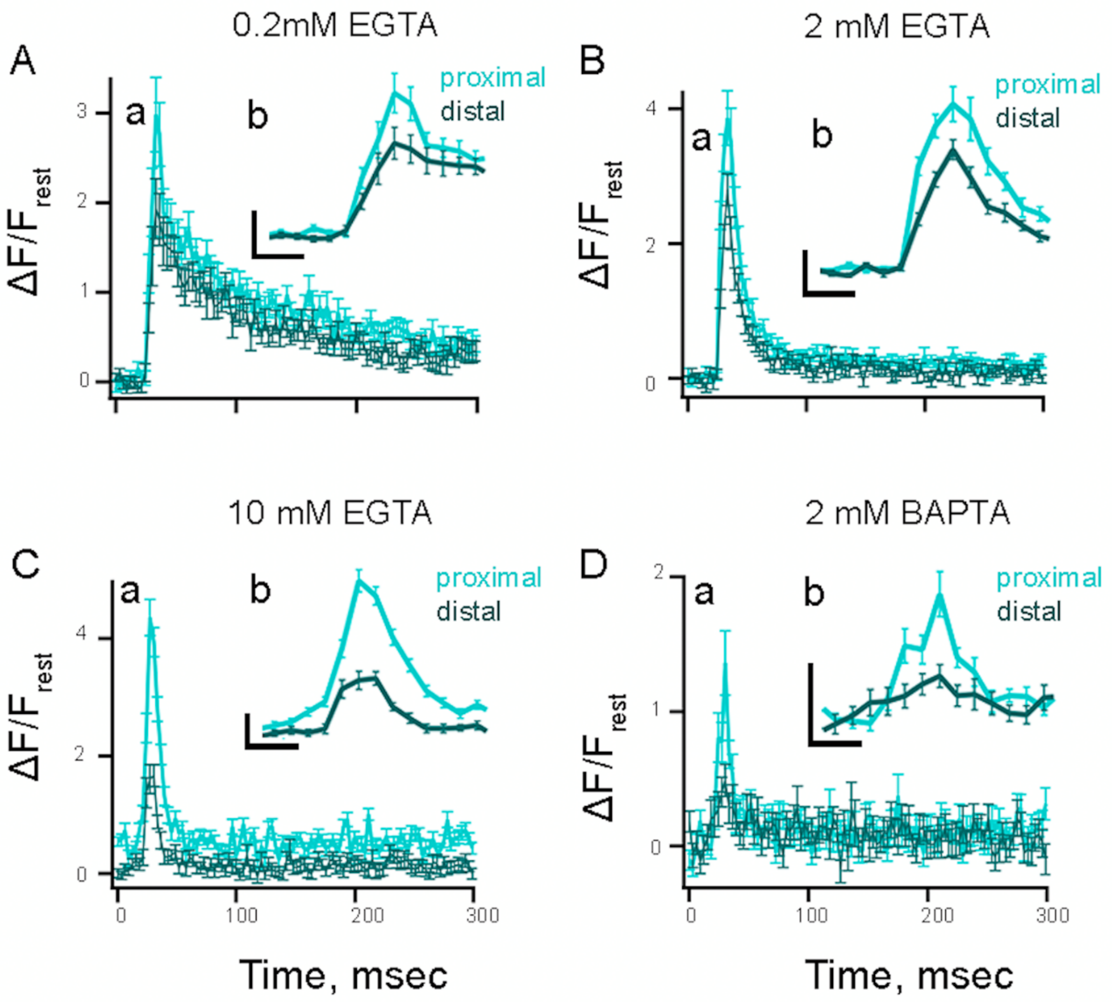


**Supplementary Fig. 3. Spatiotemporal properties of $Ca^{2+}$ microdomains along the synaptic ribbon in the RBC terminal.**

**Aa-Da.** Average temporal fluorescence intensity (normalized to $\Delta F/F_{rest}$) of proximal (light cyan), and distal (dark cyan) $Ca^{2+}$ signals measured with Cal520HA-RBP as a function of time for distinct ribbon locations with pipette solution containing (**A**) 0.2 mM EGTA, (**B**) 2 mM EGTA, (**C**) 10 mM EGTA, or (**D**) 2 mM BAPTA. Statistics comparing proximal vs. distal in each condition can be found in Supp. Table 2 below. The currents were not significantly different between conditions (Mean current amplitude in0.2mM EGTA: 49.8 ± 2.2 pA, 2mM EGTA: 53.5 ± 2.5 pA, 10 mM EGTA: 43.1 ± 2.3 pA, 2mM BAPTA: 45.6 ± 4.2 pA; 0.2mM EGTA vs. 2mM EGTA: $p = 0.29$, 0.2mM EGTA vs. 10 mM EGTA: $p = 0.059$, 0.2mM EGTA vs. 2mM BAPTA: $p = 0.39$.

**Ab-Db.** The temporal profile of events between 10-50 ms (Ab and Db) and 5-45 ms (eb and hb) was expanded for better visualization. Scale bars: vertical, 1 ($\Delta F/F_{rest}$); horizontal, 10 ms.

| | Proximal ($\Delta F/F_{rest}$) | Distal ($\Delta F/F_{rest}$) |
|---|---|---|
| 0.2 mM EGTA | 3.0 ± 0.4 (N=19) | 1.9 ± 0.3 (N=19) |
| 2 mM EGTA | 3.9 ± 0.4 (N=23) | 2.8 ± 0.2 (N=23) |
| 10 mM EGTA | 4.4 ± 0.3 (N=47) | 1.7 ± 0.2 (N=47) |
| 2 mM BAPTA | 1.4 ± 0.3 (N=22) | 0.5 ± 0.1 (N=22) |

**Supplementary Table 2**. **Effect of exogenous $Ca^{2+}$ chelators alters the $Ca^{2+}$ signal gradient along the synaptic ribbon measured with Cal520H-RBP.** There were significant differences between proximal vs distal measured as $\Delta F/F_{rest}$, in all conditions as found through paired-sample t-test analysis performed on RStudio. Differences were smaller between proximal vs distal $Ca^{2+}$ signals in 0.2 mM EGTA and 2 mM EGTA conditions, but more prominent with 10 mM EGTA, and further enhanced with 2 mM BAPTA (0.2 mM EGTA: proximal vs distal p = 0.00135, n = 19; 2 mM EGTA: proximal vs distal $p = 7.4\cdot10^{-4}$, n = 23; 10 mM EGTA: proximal vs distal $p = 1.1\cdot10^{-7}$, n = 28; 2 mM BAPTA: proximal vs distal p = 0.0013, n = 22).

| Symbol | Value / units | Description |
| --- | --- | --- |
| $C$ | μM | $Ca^{2+}$ concentration, $[Ca^{2+}]$ |
| $B$ | μM | Free (unbound) buffer concentration, [B] |
| $B^*$ | μM | Bound buffer concentration, [CaB] |
| $C_0$ | 0.1 μM | Resting background $[Ca^{2+}]$ |
| $D_C$ | 0.22 $\mu m^2$/ms | Intracellular diffusivity of $Ca^{2+}$ ions |
| $D_B$ | 0 or 0.05 $\mu m^2$/ms | Buffer diffusivity (two values used) |
| $B_{total}$ | 200 μM or 1.4 mM | Total buffer concentration (two values used) |
| $k^+$ | 0.2 $(\mu M\ ms)^{-1}$ | Buffer-$Ca^{2+}$ binding rate |
| $k^-$ | 0.4 $ms^{-1}$ | Buffer-$Ca^{2+}$ unbinding rate |
| $K_D$ | 2 μM | Buffer affinity |
| $A_{NCX}$ | 0.4 μM μm/ms | Maximal extrusion rate by NCX exchanger |
| $A_P$ | 0.03 μM μm/ms | Maximal extrusion rate by SERCA & PMCA pumps |
| $K_{NCX}$ | 1.5 μM | Affinity of NCX exchanger |
| $K_P$ | 0.3 μM | Affinity of SERCA/PMCA pumps |

**Supplementary Table 3**: Model parameters for $Ca^{2+}$ diffusion, buffering, and clearance. Simulations were performed assuming an endogenous buffer with a total concentration of either 1.4 mM (total resting buffering capacity of 720 [6-8], or a lower concentration of 200 μM corresponding to a buffering capacity of 100. Simulations in **Fig. 7** assumes immobile endogenous buffer, while **Supplementary Fig. 4** assumes a typical value of buffer mobility of 0.05 $\mu m^2$/ms. The $Ca^{2+}$ clearance parameters are adapted from [9-15]. Note that flux units of (μM μm)/ms are equivalent to $10^{-21}$ mol/($\mu m^2$ms) = 602 ions/($\mu m^2$ms). Properties of EGTA and BAPTA (not listed here) are summarized in [7].

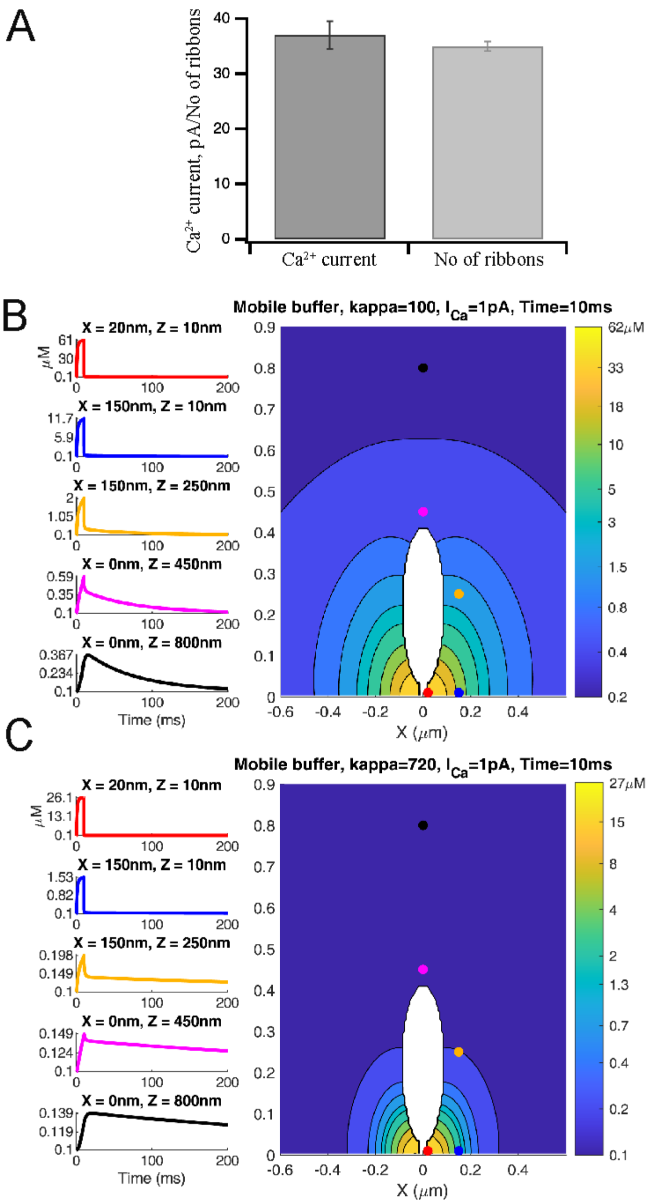


**Supplemental Fig. 4 Simulated effect of the endogenous mobile buffer of different concentrations on [$Ca^{2+}$] dynamics in response to a 10ms pulse.**

**A.** Estimation of $Ca^{2+}$ current per ribbon. The average $Ca^{2+}$ current (dark gray) and the number of synaptic ribbon fluorescence labeled with TAMRA-RBP (light gray) suggested that retinal rod bipolar ribbon synapses have a $Ca^{2+}$ current of 1pA/ribbon.

**B-C.** Simulation results are analogous to **Figure 7**, except that no exogenous buffer is included, and the endogenous buffer is mobile, with a diffusion coefficient of 0.05 μm$^2$/ms. Endogenous buffer concentrations were (**A**) 200μM (resting buffering capacity 100) or (**B**) 1.44 mM (resting buffering capacity 720).

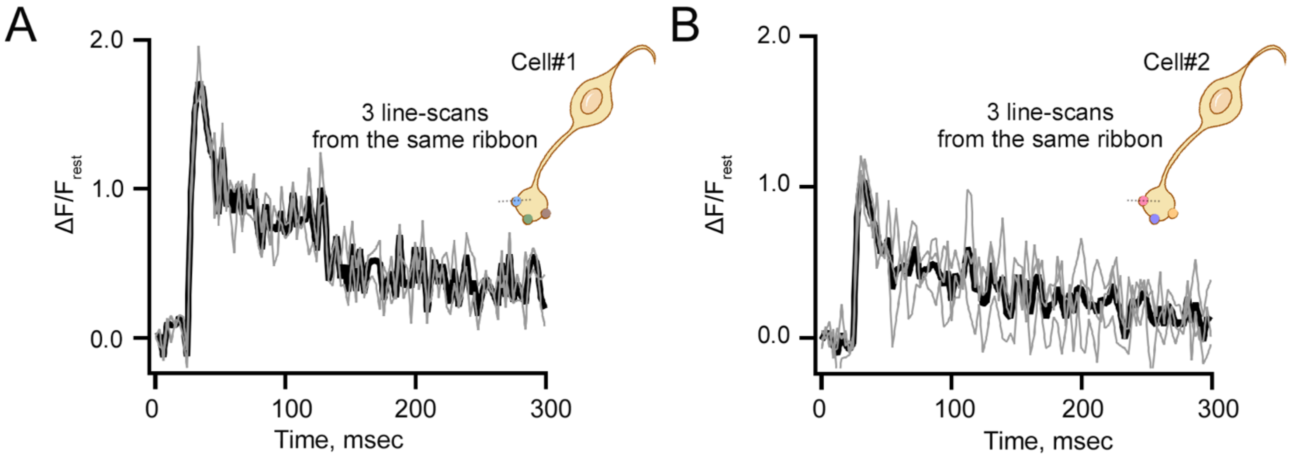


**Supplementary Fig. 5. Variability in $Ca^{2+}$ transients in response to brief stimuli.**

**A-B.** Spatially averaged Cal520HA fluorescence as a function of time at ribbon proximal location. The average (black trace) of 3 stimuli (gray traces) at a single ribbon active zone/$Ca^{2+}$ microdomain was obtained from two RBCs as described in **Fig.2A**. Note the amplitude variability between two cells (#1, panel a and #2, panel b) with similar $Ca^{2+}$ currents.

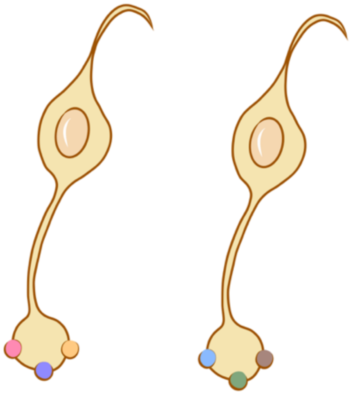

| Cell | Ribbon | Measurement Number | Data Processing | Data Presentation |
|---|---|---|---|---|
| Cell 1 | Ribbon 1 | Measurement 1 | All measurements from ribbon 1 are averaged. | Cell A |
| | | Measurement 2 | | |
| | | Measurement 3 | | |
| | Ribbon 2 | Measurement 1 | All measurements from ribbon 2 are averaged. | |
| | | Measurement 2 | | |
| | | Measurement 3 | | |
| | Ribbon 3 | Measurement 1 | All measurements from ribbon 3 are averaged. | |
| | | Measurement 2 | | |
| | | Measurement 3 | | |
| Cell 2 | Ribbon 1 | Measurement 1 | All measurements from ribbon 1 are averaged. | Cell B |
| | | Measurement 2 | | |
| | | Measurement 3 | | |
| | Ribbon 2 | Measurement 1 | All measurements from ribbon 2 are averaged. | |
| | | Measurement 2 | | |
| | | Measurement 3 | | |
| | Ribbon 3 | Measurement 1 | All measurements from ribbon 3 are averaged. | |
| | | Measurement 2 | | |
| | | Measurement 3 | | |

**Supplementary Fig. 6. Diagram of data presentation for ribbon variability between cells.**

**Top panel.** An example of two rod bipolar cells, each containing three ribbons (the first cell has ribbons depicted in pink, purple, and orange; the second cell has ribbons depicted in blue, green, and brown).

**Bottom panel.** Table explaining how the data is presented in **Fig. 8A-C**.

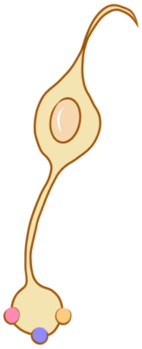

| Cell | Ribbon | Measurement Number | Data Presentation |
|---|---|---|---|
| Cell 1 | Ribbon 1 | Measurement 1 | Cell A |
| | | Measurement 2 | Cell A |
| | | Measurement 3 | Cell A |
| | Ribbon 2 | Measurement 1 | Cell B |
| | | Measurement 2 | Cell B |
| | | Measurement 3 | Cell B |
| | Ribbon 3 | Measurement 1 | Cell C |
| | | Measurement 2 | Cell C |
| | | Measurement 3 | Cell C |

**Supplementary Fig. 7. Diagram of data presentation for ribbon variability within individual cells.**

**Top panel.** Example of a rod bipolar cell containing three ribbons (depicted in pink, purple, and orange).
**Bottom panel.** Table explaining how the data is presented in **Fig. 8D-F**.

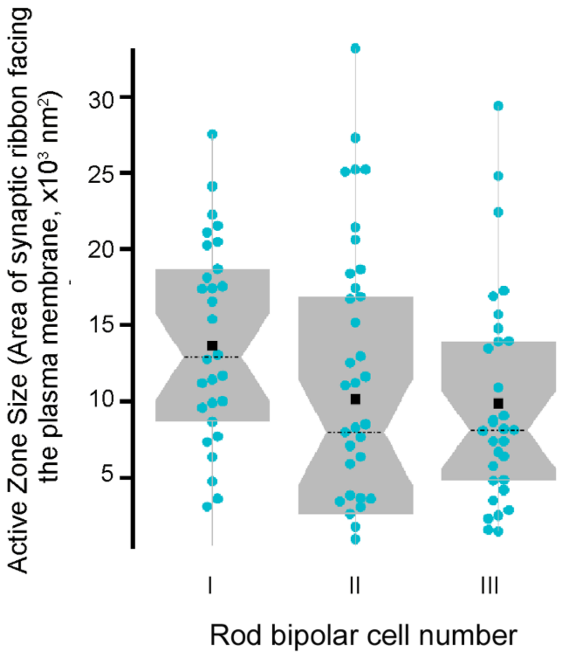


**Supplementary Fig. 8. Three RBCs active zone reconstructed from Serial block-face scanning electron microscopy.**

The box and whisker plots summarize the area of the individual ribbon associated with the plasma membrane measured in serial sections of each of the three RBCs from main **Fig. 9A**. The solid cyan circles in the box and whisker plots show the measurements of individual synaptic ribbon measurements, with the average shown as a solid black square and median values as horizontal black dotted lines. The boxes represent the 25th-75th percentiles tests. Note that RBCII contained 7 floating ribbons that were not included in main **Fig. 9A** since they were not attached to the plasma membrane.
